# Single-Cell Analysis of Microglia and Monocyte Dynamics Uncover Distinct TNF-α-driven Neuroimmune Signatures after Intracerebral Hemorrhage

**DOI:** 10.1101/2024.12.24.630187

**Authors:** Yuki Kawamura, Conor W. Johnson, Jonathan DeLong, Lucas Paulo de Lima Camillo, Sofia E. Velazquez, Munetomo Takahashi, Hannah E. Beatty, Ryan Herbert, Branden J. Cord, Charles Matouk, Michael Askenase, Lauren H. Sansing

## Abstract

Innate immune cells contribute to both secondary brain injury and repair following intracerebral hemorrhage (ICH). However, the specific signaling pathways that govern initial inflammatory and subsequent reparative myeloid programs in living patients remain poorly understood. To better characterize mononuclear phagocyte cell changes over time, we generated a single-cell transcriptomic dataset of paired hematoma clot evacuates and peripheral blood samples from 10 patients following ICH (5 - 290 hours). We identified distinct populations of activated and TNF-low microglia, as well as a unique, highly activated population of CD14^+^ monocytes in the hematoma. Perturbation analysis identified TNF signaling as the primary driver of hematoma monocyte activation. Custom temporal trajectory analysis using single-cell foundation model embeddings found that this TNF response in monocytes was transient, peaking early after hemorrhage and decreasing over the following 48 hours as monocytes shifted to reparative transcriptional programs. Transiently activated microglia emerged as the likely acute source of TNF, signaling through monocyte TNFR2. Surprisingly, acute TNF signaling in CD14^+^ monocytes was also associated with better severity-adjusted neurological outcomes both in our cohort and an independent validation cohort. These findings suggest acute TNF signaling between activated microglia and hematoma-associated monocytes, particularly through TNFR2, may contribute to recovery following ICH.

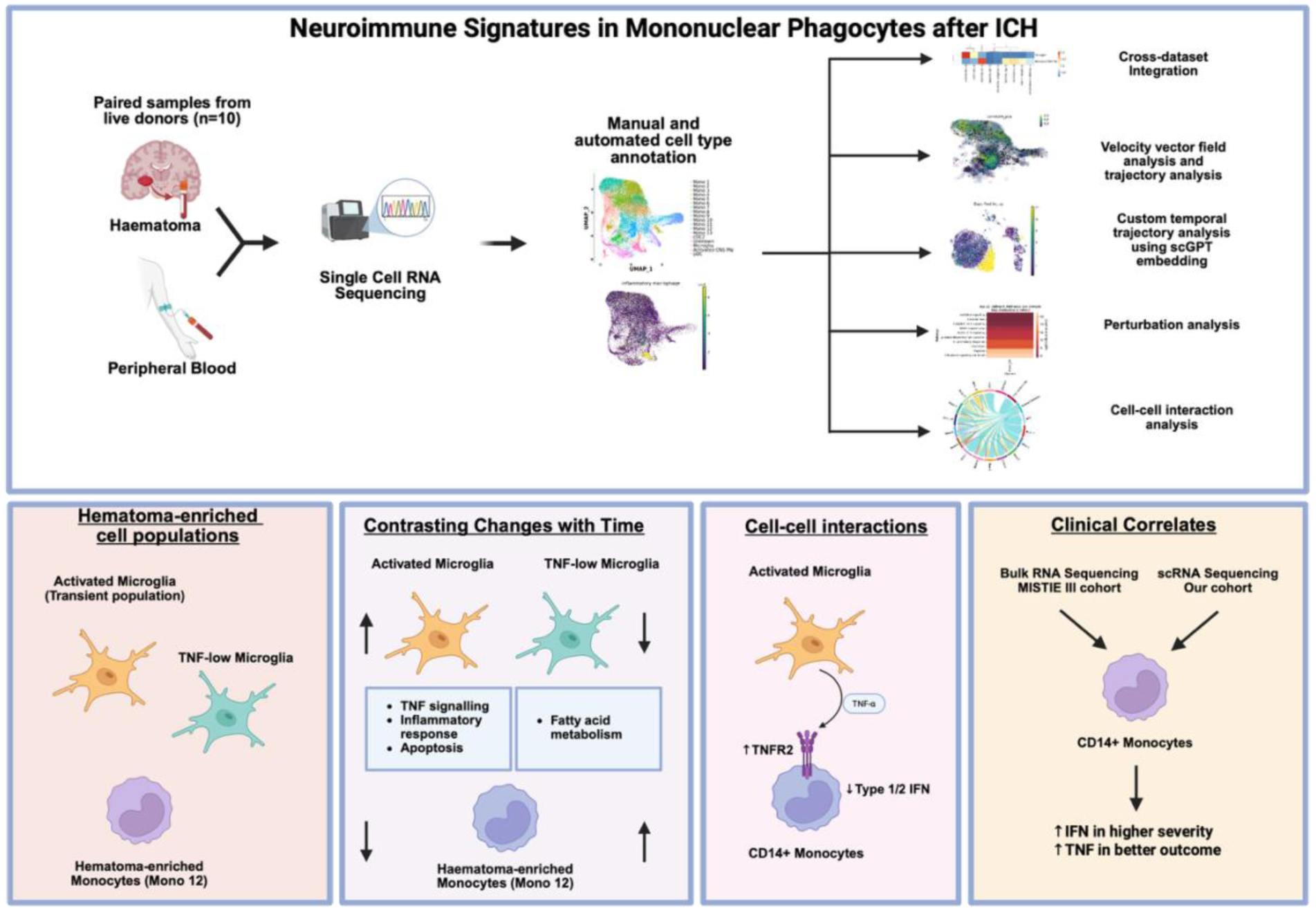

## INTRODUCTION

Intracerebral hemorrhage (ICH) comprises 27 percent of all strokes, affecting more than 3 million people worldwide annually^1^. It is the most severe form of stroke, resulting in a mortality rate of above 40% after one year and the loss of 102.2 million disability-adjusted life-years worldwide^1, 2^. Whilst current treatments focus on acute critical care interventions and rehabilitation, there are no recommended treatments that limit ongoing tissue injury following ICH. One potential target for emerging therapeutics is activated immune cells, which could improve outcome by modulating harmful immune responses and promoting tissue repair in the hemorrhagic brain.

Previous studies have demonstrated complex temporal changes in hematoma myeloid populations ^3–5^. Monocytes and microglia in particular have been implicated in contradictory roles in the hemorrhagic milieu, including aggravation of injury and inflammation, phagocytosis of erythrocytes, and neuronal repair^6^. Historically, it has been held that pro-inflammatory “M1” populations promotes a damaging early response and anti-inflammatory “M2” populations promotes a beneficial late response^7^. In this simplified view, inhibiting the M1 phenotype and polarizing cells towards the M2 phenotype would be sufficient in attenuating immune-mediated secondary injury after a brain insult. However, more recent studies have demonstrated populations which do not fit into the these simplified phenotypes^8^ as well as populations which express both M1 and M2 marker genes^9^. An example are disease-associated microglia (DAM), which exert protective effects in neurodegenerative disease through plaque phagocytosis^10^. Acute activation of myeloid phagocytic activity also appears to improve outcomes in experimental ICH by promoting hematoma clearance^11, 12^, but the role of myeloid cells in living ICH patients is poorly characterized. Despite the importance of these cell populations in the post-ICH pathophysiological response, the relative roles of brain-resident microglia and recruited monocytes has been difficult to define due to i) the difficulty of obtaining live samples from the hematoma and surrounding tissue, ii) the rapid evolution of myeloid responses during the acute stage of ICH, and iii) the complexities in defining the ontogeny of mononuclear phagocyte populations in an inflammatory setting^13^.

Herein, we present analyses of paired hematoma and peripheral blood samples from patients undergoing hematoma evacuation, which elucidate temporal changes in the monocyte and microglia populations at a single-cell resolution. These paired samples from live patients provide a rare glimpse into the complex dynamics of immune activation in the post-ICH hematoma, and the collection of peripheral blood also allows for inter-patient variability to be appropriately controlled. Using trajectory analysis of transcriptional signatures and dataset integration, we elucidate the distinct activation signatures of three mononuclear phagocyte populations in hematomas and predict their likely ontogeny as monocyte-derived vs microglia-derived. Our findings suggest the presence of a previously unappreciated and potentially beneficial TNF signaling circuit which likely originates from microglia and defines the response of hematoma CD14^+^ monocytes during the acute stage of ICH.

## RESULTS

### Single cell temporal atlas of mononuclear phagocytes in intracerebral hemorrhage

We performed single-cell RNA sequencing on paired hematoma and peripheral blood samples obtained from 10 patients undergoing hematoma clot evacuation at different timepoints following ICH (Figure 1A, Table S1). Patient age ranged from 51 to 71 years old (median: 59.5 years), with 6 males and 4 females. Time of sample collection ranged from 5.7 hours to 290.5 hours post-ICH (median: 47.2 hours). Of note, granulocytes were intentionally depleted during sample processing, as rapid degranulation during may have affected other cell activation states and prior neutrophil bulk sequencing had shown minimal outcome-associated transcriptional changes. This allowed for adequate representation of other myeloid cell types in our dataset^3^. Lymphocytes and non-myeloid CNS resident cells were also excluded from our analysis for this myeloid-focused study. After quality control and removal of doublets, we obtained 35,934 mononuclear phagocytes in peripheral blood and 7,505 mononuclear phagocytes in the hematoma. We identified 6 broad cell types in our dataset: CD14^+^ classical monocytes, CD16^+^ nonclassical monocytes, classical dendritic cells (cDC), microglia, plasmacytoid dendritic cells (pDC), and a small population of unclear identity (Figure 1B). Louvain clustering yielded 18 clusters (Figure 1C-E), which expressed varying levels of lineage marker genes (Figure S1, 2).

**Fig. 1.**
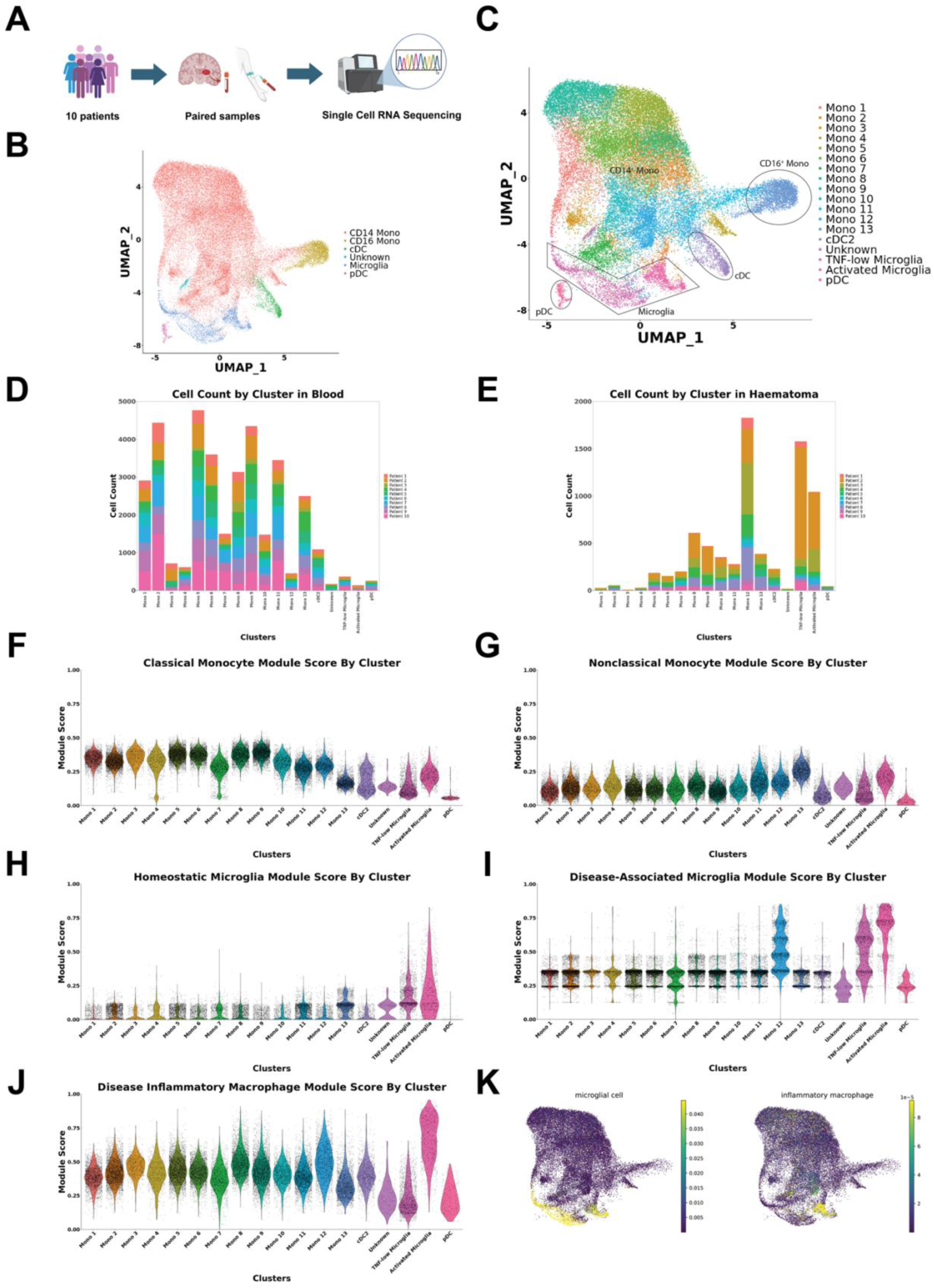
Overview of Dataset. A. Schema of experimental approach. B. UMAP plot of major cell lineages. C. UMAP plot demonstrating identified mononuclear phagocyte clusters. D, E. Cell count by cluster in blood and haematomas, respectively. F, G. Classical and non-classical monocyte module scores by cluster, respectively. H, I. Homeostatic and disease-associated microglia module score by cluster, respectively. J. Disease inflammatory macrophage module score by cluster. K. UMAP plot illustrating automated cell type annotation scores.

We identified monocyte and microglia populations in this clustering using previously established signatures for these cell types^14–16^. Twelve clusters demonstrated high expression of classical (CD14^+^) monocyte signatures (Figure 1F), whereas one cluster demonstrated high expression of non-classical (CD16^+^) monocyte signatures (Figure 1G). One CD14^+^ cluster (Mono 12) was particularly enriched in hematomas compared to blood (Figure 1E). Although microglia populations have distinct signatures under homeostatic conditions, they often downregulate their canonical markers in inflammation^10, 16^, rendering it difficult to separate them from other resident macrophage populations. We used the genes *P2RY12*, *SALL1*, *TMEM119*, *CX3CR1*, *OLFML3*, *GPR34*, *TGFBR1*, and *C1QA* to define the homeostatic microglia gene signature in our cohort (Figure 1H, S3)^15, 16^.

To better characterize the activation states of our monocyte and microglia populations, we examined their enrichment for gene signatures of DAM and disease inflammatory macrophages (DIM) associated with neurodegenerative diseases^8^. Both microglia clusters showed enrichment for the DAM signature characterized by expression of genes including *SPP1, B2M, TYROBP, TREM2, MAMDC2* (Figure 1I, S4), with stronger enrichment observed in one cluster, suggesting a potentially increased activation signature. Strikingly, we noted that this signature was also elevated in Mono 12. This cluster and one microglia cluster was also enriched for a previously described DIM signature (characterized by elevated *FOS*, *JUN*, *CCL4*, *CD83, EIF1*, etc.) seen in neurodegenerative diseases and traumatic brain injury (Figure 1J)^8^.

Given the low expression of canonical markers and overlapping enrichment for monocyte, microglia, and macrophage transcriptional signatures in hematoma mononuclear phagocytes, we independently confirmed our manual labels using a transformer-based cell type assigning algorithm trained on >22 million cells^17^. Rather than using a limited number of marker genes to define cell types, this approach can take full advantage of the high dimensionality of scRNA-seq datasets and incorporates a wider biological context. Automatic cell type annotation confirmed the presence of one cell cluster with only a microglia signature (Figure 1K), which we labelled as TNF-low microglia, and another cluster with both microglia and macrophage signatures, which we labelled as activated microglia. Taken together, we found two distinct microglia populations which have downregulated canonical markers in favor of disease-associated markers.

### Activated microglia characterize the acute phase response

CNS-resident microglia are the first non-neuronal responders following brain hemorrhage, secreting cytokines and chemokines that recruit peripheral immune cells^18^. Therefore, we next aimed to investigate the transcriptional signatures of our two microglia populations in greater depth. TNF-low microglia were characterized by expression of RPS and RPL ribosomal protein genes (*RPS2, RPL35, RPL8, RPL7A*) and HIF1A target genes (*ALDOA, GAPDH*), with lower TNF-response gene expression (Figure 2A). These genes were similar to markers of microglia populations associated with plaque phagocytosis in Alzheimer’s disease patients^19^. Indeed, top marker genes expressed in this cluster were associated with several neurological diseases on the KEGG curated gene pathway database (Figure 2B).

**Fig. 2.**
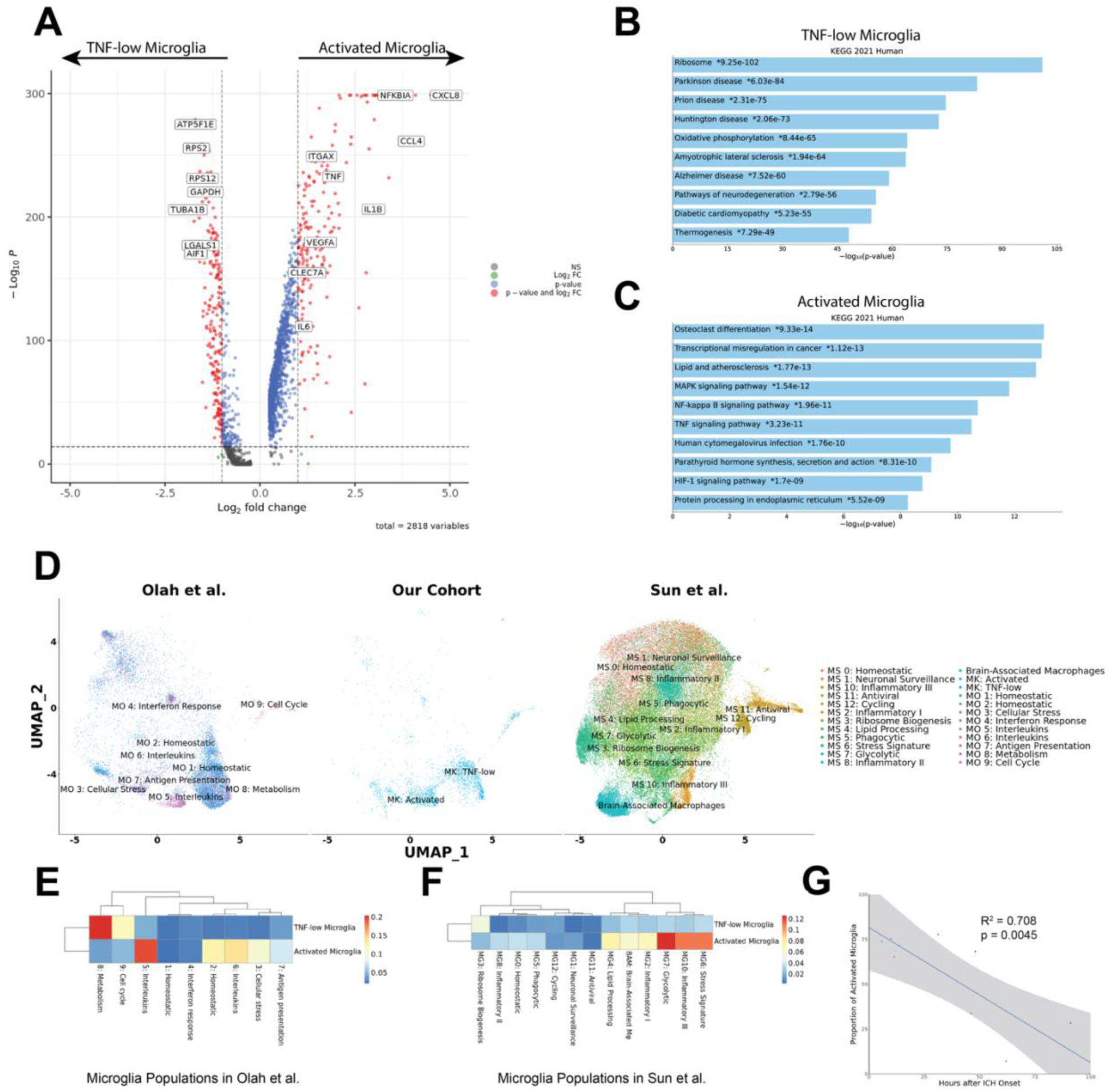
Characterisation of microglia subsets in haematomas. A. Volcano plot showing top differentially regulated genes. B, C. KEGG gene sets associated with marker genes for TNF-low microglia and activated microglia, respectively. D. Integrated UMAP plots stratified by study. E,F. Heatmap showing gene overlap score between clusters in our study and clusters in Sun et al.^20^ and Olah et al^21^. G. Streamline plots showing velocity field directionality. H. Ratio of activated microglia vs total microglia over time in the first 100 hours.

We then integrated our dataset with >200,000 cells in published microglia datasets^20, 21^ to assess whether this population corresponded to previously reported microglia populations. Both clusters overlapped with microglia from published datasets (Figure 2D, E). There was a particularly close alignment between TNF-low microglia and a disease-associated microglia subset enriched in metabolic genes and paired helical filaments-tau signatures, suggesting this population to be a conserved cross-disease responder to damage in the brain.

On the other hand, activated microglia were characterized by expression of TNF-regulated chemokine and inflammatory genes such as *CXCL8, CCL3, NFKBIA, KLF2, KLF6* (Figure 2A).

Top markers of this cluster were associated with KEGG pathways including lipid processing, MAPK, TNF, and HIF-1 pathways (Figure 2C). Activated microglia were most similar to interleukin-enriched, stress-associated, and inflammatory populations described in the literature (Figure 2E, F). Notably, the populations in the literature similar to our activated microglia were shown using scATAC-seq to be activated by HIF-1A, the canonical hypoxia transcription factor (*21*). RNA velocity analysis further demonstrated high cell transition probabilities from TNF-low to activated microglia, suggesting a potential origin of activated microglia from more stable TNF-low microglia (Figure S6D). Interestingly, this population expressed high levels of *MERTK* and *AXL*, two phosphatidylserine receptors involved in erythrocyte phagocytosis, which was associated with functional recovery after ICH in mice and humans^12^. We note that although some of the stress-associated genes elevated in activated microglia were included in a gene set associated with stress induced by sample processing mainly due to enzymatic digestion^22^, this is to be expected in the hematoma where matrix metalloproteinases are acutely elevated^23^.

Furthermore, we confirmed that our results were reproducible after selecting for cells with low expression of non-TNF-related genes in the gene set (Fig. S6)^24–26^. Hence, activated microglia appear to be a *bona fide* stress- and inflammation-related microglial population.

Finally, the proportion of activated microglia among total microglia decreased with time after hematoma onset (Figure 2G), becoming nearly absent after 100 hours. In summary, microglial dynamics following ICH is defined by two clusters: an acute, inflammatory population of activated microglia, and a subacute, metabolically active population of TNF-low microglia associated with neurodegenerative disease.

### Hematoma monocytes skew towards TNF-a signaling and away from interferon signaling

We then sought to characterize the CD14^+^ monocyte populations in the peripheral blood and hematomas. Populations dramatically enriched in peripheral blood included the Mono 3 cluster, characterized by expression of metallothionein genes such as *MT1E*, *MT2A, MT1X*, as well as the Mono 4 cluster, characterized by interferon-response genes such as *ISG15*, *MX1*, *IFIT3*, *IFI44*, *IFI44L* (Figure 3A, S7). We found 9 populations (Mono 1-2, 5-11) which were present in both tissue compartments (“shared monocytes”).

**Fig. 3.**
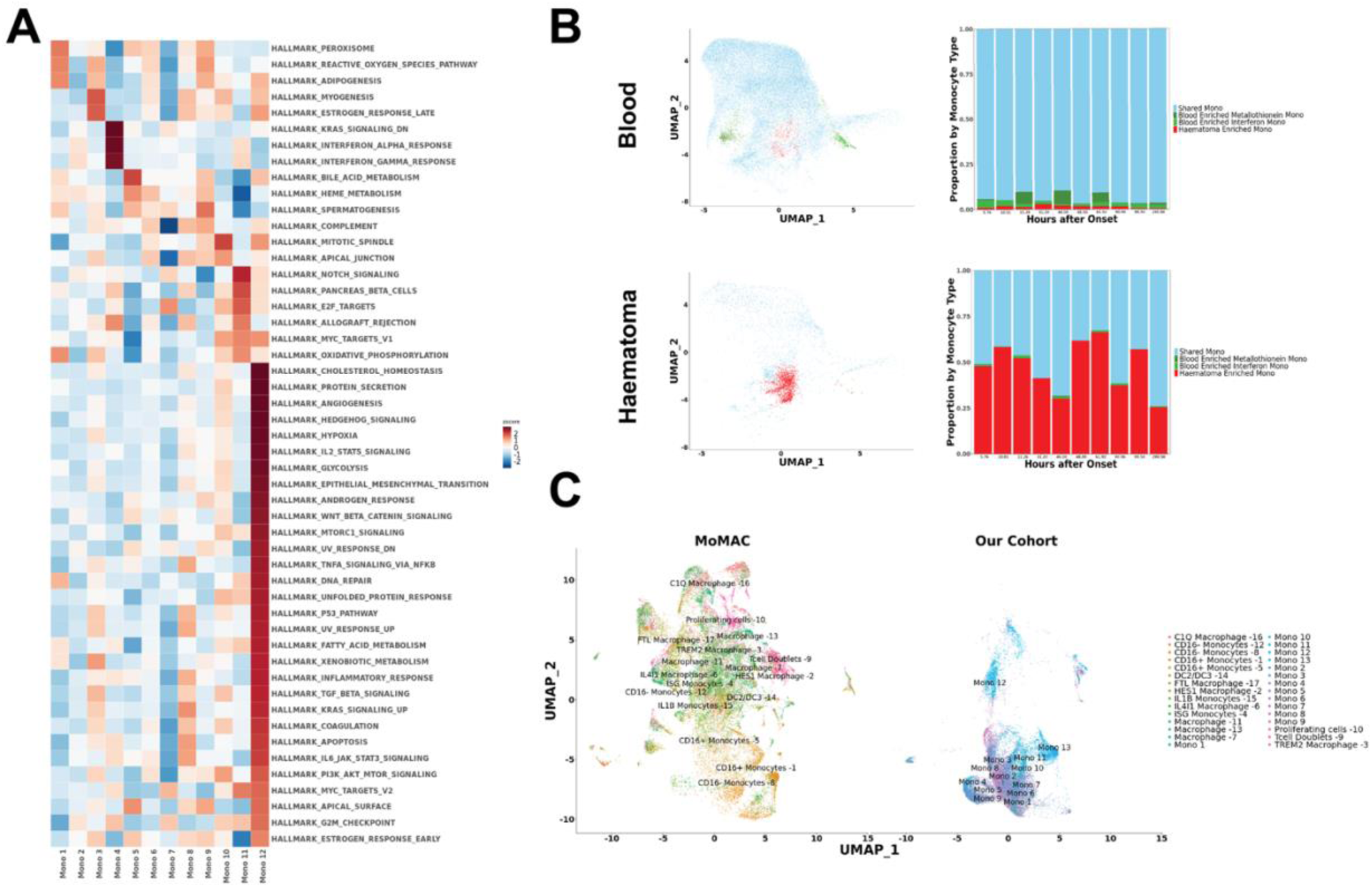
Characterisation of monocyte populations. A. Heatmap showing pathways enriched in each cluster, separated by tissue. B. UMAP plots and cell proportion plots ordered by hours after onset for blood (top) and haematomas (bottom). C. Integrated UMAP of monocytes in our dataset and a cross-tissue monocyte/microglia atlas^27^.

One monocyte population, Mono 12, was drastically enriched in hematomas (Figure 1D, E, 3C). Mono 12 cells were present in all patients (Figure S7C) and were present from as early as 5.8 hours after ICH (Figure 3B). This population was characterized by high expression of genes co-regulated with the TNF pathway such as *CCL2*, *CXCL2*, and *CXCL3* (Figures 3A, S1). In addition, this population was enriched in pathways such as hypoxia, glycolysis, angiogenesis, TNF, and had low expression of interferon-associated genes (Figure 3A, S1). Comparing our monocyte dataset with a cross-tissue monocyte/macrophage atlas combining 41 datasets^27^ demonstrated that the Mono 12 corresponded to inflammatory “IL1B Monocytes”, a cell population expanded in patients with severe COVID-19 and tumors (Figure S7B).

Overall, hematoma and peripheral blood monocytes shared some similar characteristics, but with a key difference of hematoma monocyte polarization towards TNF signaling and away from type 1 interferon signaling.

### Mono 12 is associated with a validated hematoma signature

Given our smaller sample size, we next sought to corroborate the upregulation of the Mono 12 transcriptional signature in hematoma using a larger, independent cohort. We compared our CD14^+^ monocytes with previously published CD14^+^ monocyte bulk sequencing data that was generated as a part of ICHseq substudy of the MISTIE III trial (Figure 4A)^3^. In that study, matched hematoma and peripheral blood samples were collected from ICH patients daily as a part of catheter hematoma effluent drainage. We only included samples collected in the first 100 hours post-ICH in this analysis to time-match our scRNA-seq cohort (hematoma: 33 samples from 19 patients, blood: 43 samples from 21 patients).

**Fig. 4.**
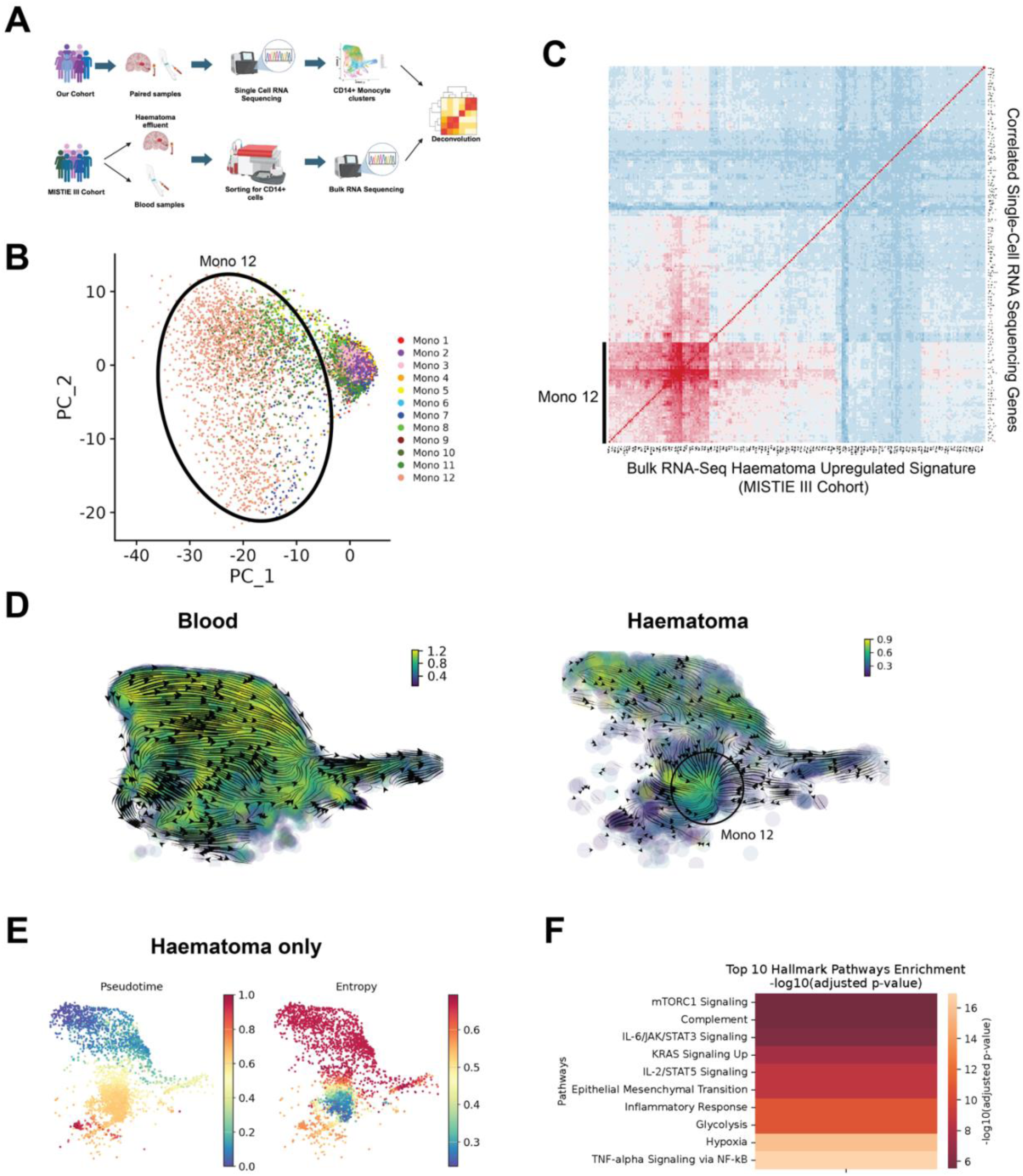
Characterisation of haematoma-enriched Mono 12. A. Schema of validation of haematoma signatures. B. Principal components plot of CD14+ monocytes based on top 250 genes enriched in haematoma compared to peripheral blood in bulk sequenced samples from the MISTIE III cohort. C. Pairwise correlation matrix of bulk RNA-Seq haematoma-associated genes (columns) and top correlated genes in scRNA-Seq dataset (rows). D. Streamline plots showing velocity field vector directions and coloured by vector field curvature. E. Pseudotime and entropy plots for CD14+ monocytes in haematomas. F. Heatmap of pathways associated with perturbation from shared monocyte populations to haematoma-enriched monocytes (Mono 12).

Differential expression analysis between hematoma and peripheral blood and subsequent PCA of the top 250 hematoma-enriched genes revealed a clear separation of Mono 12 from the rest of the monocyte clusters (Figure 4B), suggesting that Mono 12 distinctly expressed hematoma-enriched genes. We then deconvoluted bulk sequencing data by performing pairwise gene correlations with the top 250 hematoma-enriched genes for each cell and performing subsequent hierarchical clustering (Figure 4C). This demonstrated that the genes highly correlated with the hematoma signature were markers of Mono 12, validating the presence of this TNF-responsive hematoma monocyte population in an independent cohort of patients.

### Mono 12 cluster is associated with end stage of monocyte cell transitions

Noting the changes in monocyte populations observed in hematomas and in peripheral blood, we hypothesized that activating signals in the local milieu were responsible for the differentiation of circulating CD14^+^ monocytes into the Mono 12 cluster we observed in the hematoma. To examine this hypothesis, we leveraged RNA velocity analysis in monocytes to identify potential trajectories of differentiation across peripheral blood and upon hematoma entry based on their splicing dynamics. Although subject to methodological limitations, it can be useful for inferring cell state transitions if interpreted in the appropriate context of known biological changes.

RNA velocity analysis of monocyte populations across both tissues demonstrated a trajectory towards CD16^+^ monocytes in the blood, in agreement with previous reports of a linear differentiation path from CD14^+^ to CD16^+^ monocytes^28^. However, an additional trajectory towards Mono 12 was identified only in hematomas (Figure 4D). Mono 12 was also associated with late pseudotime and low potential (entropy) for differentiation into other cell states, suggestive of a differentiation endpoint (Figure 4E). Random walk analysis on RNA velocity vectors likewise indicated that the Mono 12 cluster was a unique potential endpoint of cell type transitions in the hematoma samples, but not in peripheral blood (Figure S8). This agrees with biological intuition, because extravasated monocytes in hematomas should initially reflect their counterparts in peripheral blood before transitioning to a unique hematoma-enriched phenotype.

In addition, we conducted an analysis of the properties of the RNA velocity vector field to identify where the greatest changes in cell transition states were occurring. Interestingly, RNA velocity vector field curvature (local rate of change) was uniform in blood regardless of cluster but was high in Mono 12 in hematomas (Figure 4D). This suggests that Mono 12 has an especially higher rate of intra-cluster state transitions, likely due to a high variation in its transcriptional signature. Taken together, these findings supported the Mono 12 cluster being a heterogenous end state of monocyte transitions in hematomas.

### TNF signaling is a major driver of perturbation in CD14^+^ monocytes

Next, we sought to identify what genes were responsible for driving transitions of monocytes towards Mono 12. We therefore employed perturbation analysis to perform causal inference^29^ on genes that were responsible for driving shifts in distribution between shared monocytes (Monos 1-2, 5-11) and hematoma-enriched monocytes (Mono 12). Our analyses inferred that the TNF and hypoxia pathways were the most important drivers of the shift in gene signatures from shared monocytes to hematoma-enriched monocytes (Figure 4F), which agreed with *in silico* predictions of hematoma signature upstream regulators in the MISTIE III samples^3^.

### Foundation model embedding captures temporal and tissue patterns without metadata

Having characterized population shifts responsible for differences by tissue compartment, we then sought to characterize how cell populations shift over time. Generative pre-trained transformers (GPT) are a type of large language models, a subset of machine learning models. Their ability to capture latent dimensions of data and to assign attention scores to identify pertinent features important in prediction makes them a useful tool in identifying underlying structure in high-dimensional datasets such as ours. Our goal was to dissect underlying changes which were associated with temporal progression in the monocyte and microglia populations.

For this purpose, we employed scGPT^30^, a GPT model pre-trained on >33 million cells to generate cell embeddings (lower-dimensional representations) in completely unsupervised fashion. We hypothesised that the diverse training data used to develop this would allow it to capture biologically meaningful signatures within ICH mononuclear phagocytes. scGPT embedding was performed solely using the gene expression matrix — metadata such as time and tissue compartment were not provided to the model.

scGPT embedding clearly separated peripheral blood from hematoma samples and monocytes from both TNF-low and activated microglia (Figure 5A, B). These results suggested that the embeddings effectively captured the different transcriptional signatures in our dataset and prompted us to explore temporal relationships.

**Fig. 5.**
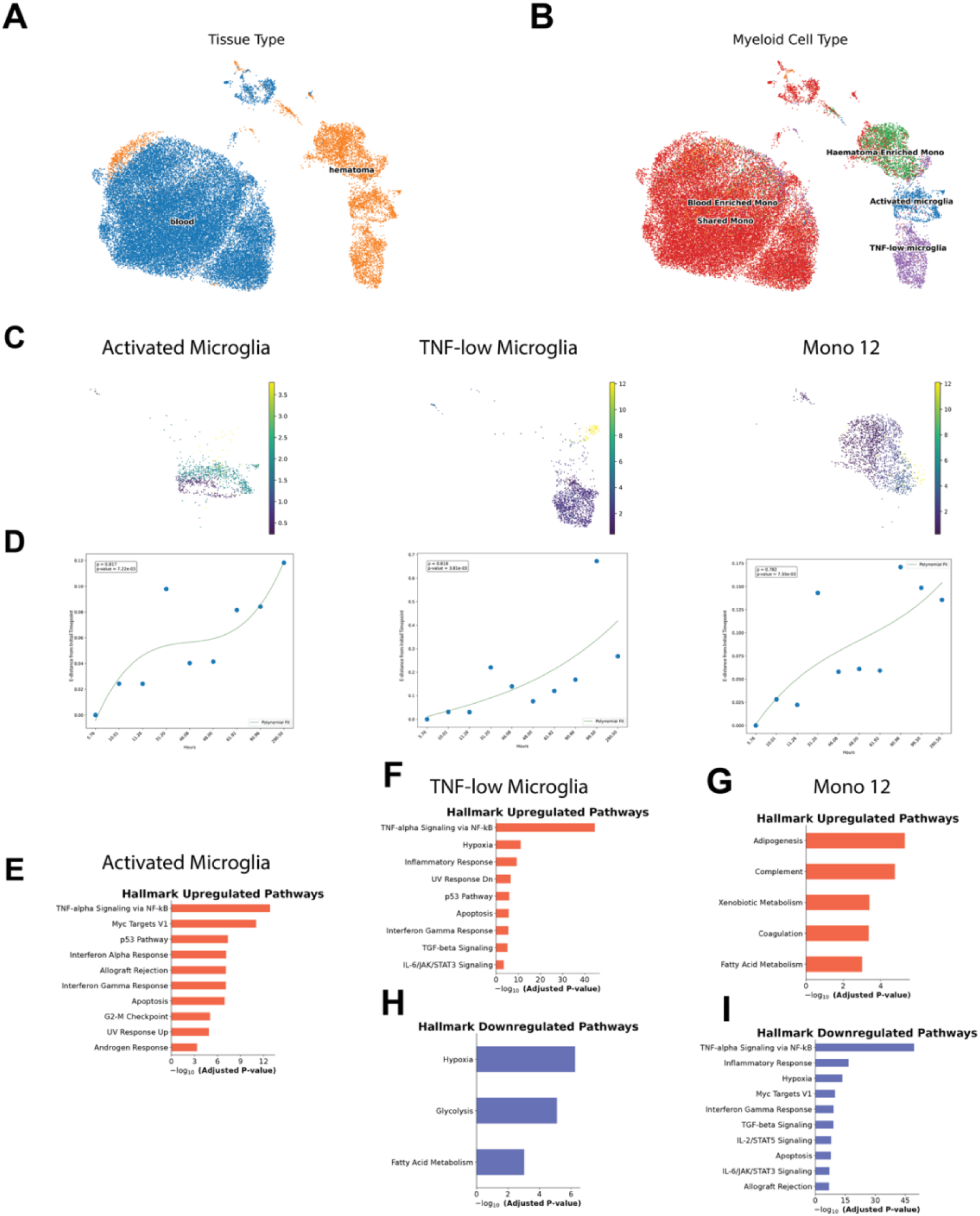
Temporal changes in the dataset captured by scGPT model. A. UMAP of scGPT feature embedding coloured by tissue and cell type, respectively. C. UMAP of each cluster coloured by days post ICH. D. Energy distance from initial time point for activated microglia, TNF-low microglia, and Mono 12 in haematomas. E-G. Gene sets upregulated with embedding distance in activated microglia, TNF-low microglia, and mono 12, respectively. H,I. Gene sets downregulated with embedding distance in TNF-low microglia and Mono 12, respectively

When stratified by sample timepoint, the embedding space demonstrated a sequential trend in hematoma but not blood samples (Figure 5C). We then quantified these changes by comparing the energy distance (a measure of difference in statistical distributions) per sample at each timepoint after ICH. If the embedding faithfully captured the temporal sequence, we would expect to see a correlation between the time after ICH and energy distance from cells at the earliest timepoint. Indeed, our results confirmed that embedding distance correlated with time in hematomas for most clusters — including Mono 12, TNF-low microglia, and activated microglia (Figure 5C, D). In contrast, no such correlation was observed in blood (Figure S9). This independently confirmed underlying changes with time in hematomas, but not peripheral blood, in the expression patterns of microglia and monocytes.

### Microglia and Mono 12 exhibit contrasting temporal patterns

We next analysed gene sets associated with temporal changes within major populations in hematomas using a custom temporal trajectory based on scGPT embeddings (see Methods). Both microglia populations upregulated the TNF and interferon gamma pathways with time, whereas the TNF-low microglia population downregulated the hypoxia and glycolysis pathways (Figure 5E, F, H). The activated microglia population did not have any significantly downregulated pathways over time. On the contrary, the Mono 12 cluster downregulated the TNF, inflammatory response, hypoxia, and interferon gamma pathways over time in favor of adipogenesis, complement, coagulation, and fatty acid metabolism pathways (Figure 5G, I). These upregulated pathways are consistent with tissue repair responses^31^. Importantly, we note that these findings agree with bulk RNA sequencing results in the larger MISTIE III cohort^3^. This suggested that the Mono 12 cluster appeared to mount a rapid inflammatory response and reduce the expression of these pathways shortly thereafter. We confirmed this rapid peak in TNF response by inferring the splicing rate of genes associated with the TNF pathway in the Mono 12 cluster. The transcription of many of these genes rapidly ceased after the first 48 hours (Figure S10), explaining its high velocity field curvature (Fig. 4D). The proportion of activated microglia was halved also after 48 hours (Figure 2G), suggesting a potential interplay between the cell populations.

Thus, our temporal trajectory analysis demonstrated that while TNF-associated transcriptional signatures became augmented over time in microglia, by contrast, TNF signaling was downregulated in the hematoma by monocytes in favor of pathways associated with tissue repair responses.

### Optimum transport analysis on Mono 12 cluster confirms temporal changes in TNF

We also used optimum transport — which infers cell state transitions by calculating the most parsimonious shifts in cell state distributions over time — to independently recapitulate this rapid decrease in TNF signaling in the Mono 12 cluster. Our goal was to confirm that temporal patterns inferred from gene expression embeddings could be reproduced with an alternative method that uses time as an input variable.

In this analysis, cells in the Mono 12 cluster also transitioned to cells with lower expression of the TNF over time (Figure S11A-C). Mono 12 cells also appeared to be downregulating pathways which define their cellular phenotype with time (Fig 3A, S11 B-C). In summary, the Mono 12 TNF response rapidly decreased in the first 100 hours by both optimum transport and custom scGPT temporal embedding, resulting in a loss of cluster-defining signatures early after ICH.

### Activated microglia are a master regulator of TNF signaling

In light of the observed correlation between activated microglia proportion and acute Mono 12 TNF signature, as well as previous literature that establishes microglia as the dominant post-injury producers of TNF in the CNS^32^, we hypothesized that microglial cell-cell communication could account for the observed rapid changes in monocyte TNF signaling. We employed cell-cell communication analyses using CellChat v2^33^ in monocytes and microglia in hematoma samples to address this question, which demonstrated that the TNF pathway was amongst the most significant signaling pathways between microglia and monocytes (Figure S12).

Focusing on the TNF pathway, activated microglia were found to be a major source of TNF signaling, in particular as the producer of TNF ligand (Figure 6A, B). The most highly regulated ligand-receptor pair was *TNF-TNFRSF1B* (coding for TNF Receptor 2), followed by *TNF-TNFRSF1A* (coding for TNF Receptor 1) (Figure 6C). Whilst TNFR1 is associated with classic NFkB activation and apoptosis^34^, TNFR2 is associated with anti-inflammatory signaling and alternative NFkB activation^35^. The ratio between the gene expression of *TNFR2* and *TNFR1* in CD14^+^ monocytes was significantly increased (3.1-fold; p=0.02) in hematoma samples compared to blood (Figure 3D), indicating a shift towards TNFR2 signaling *in situ*. The significant increase in *TNFR2*/*TNFR1* ratio in hematomas was also replicated in the MISTIE III cohort (Figure S13A). We also confirmed an increase in the TNFR2 downstream signaling signature in CD14^+^ monocytes (Figure S13B, C). Taken together, these results suggest that microglial-monocyte signaling, especially TNFR2 signaling, plays a critical role in the acute hemorrhagic environment. We speculate that this intercellular communication via TNF is responsible for the drastic yet transient presence of TNF response pathways observed in hematoma monocytes.

**Fig. 6.**
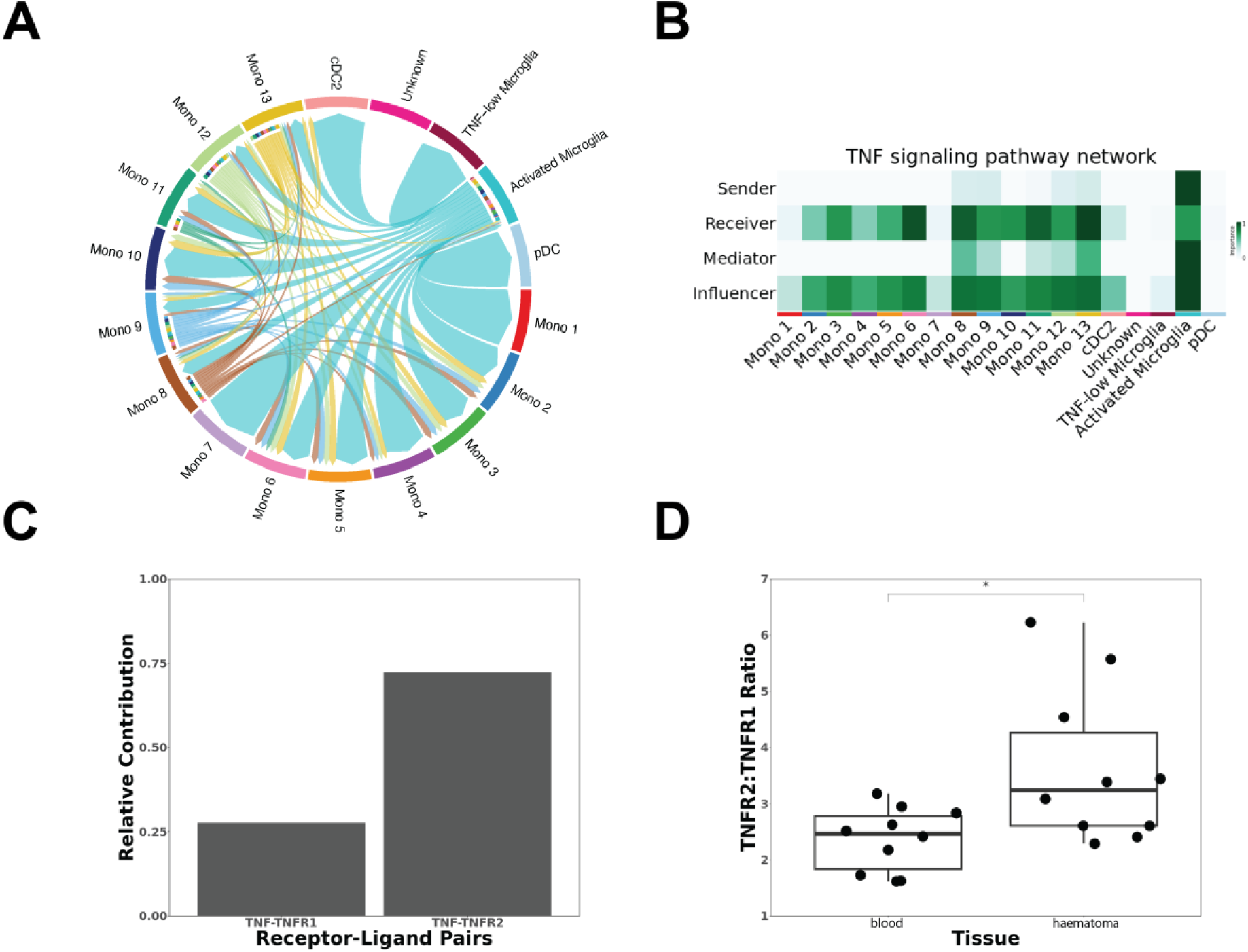
Cell-cell Interaction Analysis. A. Chord plot of TNF ligand-receptor interactions between different cell types. B. Heatmap showing different roles of various clusters in TNF signalling in haematomas. C. Relative contribution of TNF receptor-ligand pairs between microglia and monocytes. D. Relative expression of *TNFR1* and *TNFR2* in CD14^+^ monocytes. *p < 0.05, Wilcoxon test

### TNF signaling associates with outcome but not severity in CD14^+^ monocytes

We then assessed whether CD14^+^ monocyte transcriptional signatures were associated with clinical severity or outcome, in both our patient cohort and the MISTIE III cohort. We first correlated the transcriptional signature with two measures of ICH severity on presentation to hospital: large hematoma volume and low consciousness level, measured by the Glasgow Coma Scale (GCS). Whilst TNF signaling was not consistently associated with initial severity, interferon pathways were consistently associated with increased severity (Figure 7A-D, S14).

**Fig. 7.**
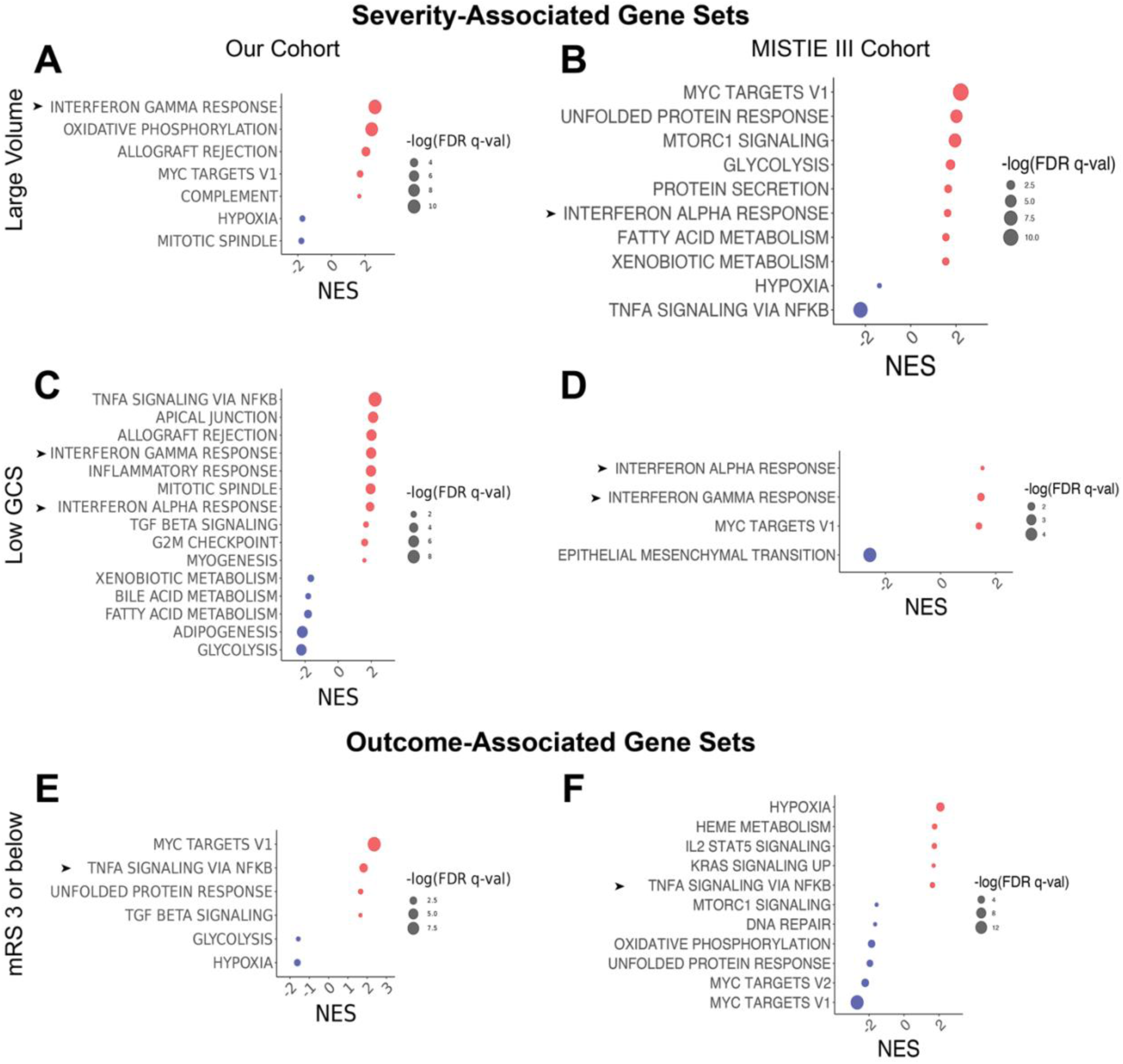
Gene sets associated with ICH severity and outcome in CD14^+^ monocytes. A, B. Gene sets enriched in hematoma samples in patients with higher pre-excavation ICH volume in our cohort and the MISTIE III cohort, respectively. C, D. Same as A and B, but gene sets associated with low GCS on presentation. E, F. Gene sets enriched in haematoma samples compared to peripheral blood in patients with better outcome, adjusted for ICH score in our cohort and the MISTIE III cohort, respectively.

Good outcome was defined as 90-day modified Rankin Scale (mRS) scores between 0 and 3 (lower scores denote less disability after stroke). We used the ICH score (which includes GCS, ICH volume, age, intraventricular hemorrhage (IVH) and ICH location as components), a well-validated clinical score for risk stratification^36^, as a covariate to avoid confounding by initial severity and demographics. The small sample size of the singe-cell sequencing cohort made several gene set correlations susceptible to outlier effects, leading to some inconsistencies across cohorts. Nevertheless, TNF signaling was associated with better outcome in both cohorts (Figure 7E, F, S15). Results in peripheral blood showed some overlap with hematoma signatures but were inconclusive (Figure S14). Taken together, these correlations with clinical variables suggest that different inflammatory pathways play distinct roles in the hematoma milieu and that, surprisingly, the microglia-monocyte TNF signaling axis in the hematoma could be beneficial after ICH, perhaps through increased TNFR2 signaling.

## DISCUSSION

This study used single cell RNA sequencing to investigate the mononuclear phagocyte response in samples of live hematoma tissue and matched peripheral blood following ICH. We generated a temporal atlas of 35,934 peripheral blood and 7,505 hematoma mononuclear phagocytes from 10 patients, characterizing the transcriptional profiles of monocytes, microglia, and dendritic cells. This dataset serves as a resource for the scientific community to explore cellular dynamics and test hypotheses regarding immune responses in human ICH.

Within monocyte and microglial populations, TNF signaling emerged as a key early transcriptional driver of multiple hematoma clusters, and such signaling in monocytes was associated with improved neurological outcomes. We identified a short-lived activated microglia population which exhibited transcriptional similarities with previously described stress- and inflammation-associated populations and was the major driver of TNF signaling in the hematoma milieu. Amongst the monocytes, Mono 12 emerged as an acute responder population that was dramatically enriched in hematomas and drove much of the hematoma signature. This cluster was defined by expression of TNF-related genes in the first 48 hours, which were then downregulated in favor of pathways associated with tissue repair responses. Total brain TNF protein peaks around 2-4 hours post-ICH in both autologous blood^37^ and collagenase^38^ rodent models, while levels of TNF and associated proteins surrounding the hematoma peak on day 3 post-ICH^39^. This timeline of activation generally aligns with the transient upregulation of TNF signaling seen in the activated microglia and Mono 12 clusters in our dataset. Our analysis suggests that the rapid loss of activated microglia-derived TNF could account for the loss of the hematoma-defining signature in Mono 12. Further studies investigating the role of this distinct monocyte population in the context of ICH recovery are warranted.

TNF signaling has been implicated in a wide range of neuroinflammatory processes, with differing effects depending on the ligand, receptor, and disease pathology. TNF protein is first produced as a transmembrane form (tmTNF) which acts through cell-cell contact, and is subsequently cleaved to a soluble cytokine (solTNF)^40^. solTNF activates TNFR1, while tmTNF can activate both receptors but functionally prefers TNFR2^41^. TNFR1 contains a death domain (DD) and is associated with classical NFkB signaling and apoptosis. On the other hand, TNFR2 does not contain a DD and activates both classical and alternative NFkB complexes, as well as some anti-inflammatory cytokines^35^. *TNFR1* is ubiquitously expressed across cell types, while *TNFR2* expression is limited mostly to immune cells^32^, and restricted to glia and infiltrating macrophages in the peri-infarct area in human ischemic stroke^42^. Plasma levels of both TNF receptors are enriched in patients who later go on to develop ICH and are associated with increased risk of death and poor functional outcome^43^.

Preclinical rodent studies have largely found TNF to play a deleterious role in ICH, which runs counter to the outcome data described herein. This may be a species difference, as we used solely human data, or due to mixed effects of TNFR1 and TNFR2. Antagonism of TNF through intracerebral injection of an antisense oligonucleotide^44^, intravenous injection of a TNF antibody^38^, and TNFR1 antagonism^45^ reduced edema volume, although reports on neurobehavioral scores have been mixed^39, 46^. Genetic knockout of TNF and TLR4 similarly decreased edema and improved survival and neurobehavioral outcomes^37, 39^. However, the limited functional outcome measures in these studies make it challenging to draw definitive conclusions about the role of TNF. Of note, TNF inhibition was not associated with lower hematoma volume at either 24^38^ or 72^46^ hours, and neurobehavioural differences were not tested past three days in most studies. One consistent finding was acute perihematomal edema reduction, but this is only weakly associated with long-term ICH outcome in humans^47^. These studies also leave unanswered questions on how the complexities of TNF activity might manifest in ICH, since they focus on soluble TNF (solTNF) and TNFR1. The authors are not aware of studies that have causally tested the effect of transmembrane TNF (tmTNF) or its primary receptor TNFR2 in murine ICH models.

Studies using models of ischemic stroke using middle cerebral artery occlusion (MCAO), however, have investigated TNF ligand and receptor forms more thoroughly. Broad inhibition of TNF via antibody injection^48^ and systemic administration of etanercept (a combined TNF inhibitor)^45^ improved outcomes, whereas intracerebroventricular injection of TNF worsened outcomes^47^. solTNF-specific inhibition was protective^47^ but both global and neuronal TNFR1 knockout were detrimental^49, 50^. Microglial-specific and global *TNFR2* knockouts had mixed effects on functional tests and other outcome measures^49–51^, but direct TNFR2 agonists have demonstrated transient functional improvement with systemic treatment^52^, and selective tmTNF expression was protective^53^. Taken together, these studies suggest solTNF has deleterious function in ischemic stroke, but that tmTNF signaling through both TNF receptors may exert neuroprotective effects. Similar neuroprotective effects were seen in a multiple sclerosis rodent model, where selective solTNF inhibition, but not combined inhibition, was neuroprotective and promoted remyelination^54, 55^, likely due to critical roles of TNFR2 signaling in microglia and neurons for remyelination^56–58^.

Although the pathophysiology of ICH, ischemia, and peripherally mediated demyelination only partially overlap, similarities in the effects of altered TNF signaling pathways suggest a possible conserved CNS damage response, accounting for its prominent role in our dataset. Our findings highlight crosstalk between activated microglia and CD14^+^ monocytes primarily through TNF acting on TNFR2, which was enriched in hematomas in both our and the MISTIE III cohort. tmTNF and TNFR2 signaling represent a potential protective mechanism of TNF in the post-ICH brain. Further investigations into the role of tmTNF and TNFR2 in ICH could help to elucidate these pathways.

Our study has several limitations. Given that hematoma evacuation is generally only performed as part of relevant clinical trials or in response to clinical deterioration, our sample size was inevitably limited. We sought to address this by validating gene signatures in an independent cohort of patients who were recruited separately and analyzed using a different method, which confirmed our main findings. Additionally, the analysis of functional outcomes in our cohort and the MISTIE III cohort could be confounded by surgical evacuation. However, we note that surgical evacuation itself did not affect outcome in the trial^59^, and we therefore believe that this confounding factor is minimal. Next, our mononuclear phagocyte-focused study intentionally excluded granulocytes as a part of tissue processing, as prior bulk sequencing work found minimal outcome-associated changes in this population^3^. Therefore, we cannot make conclusions about the role of this population in our dataset. We also did not identify any clear border-associated macrophage population in our dataset; these may present around the hematoma but were either unintentionally removed as a part of processing or were not transcriptionally distinguishable from other tissue-resident macrophage CNS populations in this inflamed, hypoxic environment. Finally, given that TNF signaling is associated with increased brain edema^37, 38^, it is possible that our study artificially minimizes harmful TNF-driven edema through evacuation, revealing protective acute phase effects. Therefore, additional studies are needed to validate the association of TNF and other pathways with functional outcome. Our findings still underline, however, that the role of TNF signaling in the acute response to ICH is likely to be multifaceted and that non-specific inhibition might be detrimental.

Overall, our study provides detailed insights into the temporal pattern of phenotypic shifts in microglia and monocytes in the brain, revealing a transient TNF-driven phenotype that is associated with better severity-adjusted outcome in our cohort and the MISTIE III cohort. Our findings establish complex, acute shifts in the hematoma immune environment, informing treatment windows and molecular targets for future immunomodulatory interventions. Our findings also provide a novel resource for researchers to test hypotheses of innate immune activation and modulation in the live hemorrhagic brain, as well as neuroinflammatory responses more broadly in living patients.

## MATERIALS AND METHODS

### Sex as a Biological Variable

Our study included both male and female human subjects.

### Study design

This study used scRNA-seq to characterize longitudinal myeloid transcriptional responses in living brain and blood following ICH, enabling unbiased identification of key cell populations and pathways. Evacuated hematoma and peripheral blood samples were collected from 10 ICH patients admitted to Yale-New Haven Hospital from March 2019 to December 2021. Surgery was performed when the clinical team determined surgical evacuation was medically necessary or if the patient was enrolled in a clinical trial evaluating the efficacy of surgical evacuation.

Hence, no randomization was performed. Hematomas were surgically evacuated using minimally invasive surgery (n=5) or craniotomy/craniectomy (n=5) (Table S1). The research was approved by the IRB of Yale School of Medicine (HIC #1506016023). All subjects or their legally authorized representative provided informed consent.

Initial target sample size was 10-15 patients, to balance robust analysis with logistical and financial feasibility. No formal power analysis was performed, as the primary focus was exploratory. All 20 collected samples were included in the final dataset – exclusion criteria were determined at the single-cell level in accordance with standard preprocessing (see supplement). One patient underwent hematoma sampling much later (290 hours) than the rest of the dataset; this was due to rapid clinical deterioration. Sampling and technical replicates within patients were not possible due to the unique sample collection circumstances.

Patient outcomes were assessed by blinded investigators using modified Rankin scale (mRS) scores 3 months post-ICH. MISTIE III patient outcomes were assessed as previously described^59^. All investigators were blinded to patient data, including severity and outcome data, for all procedures and analyses where it was not directly required.

### Cell-cell interaction analysis

Cell-cell interaction analysis was performed using CellChat2 ^33^ after filtering for non-protein interactions and subsetting for microglia clusters in haematoma tissue.

### Velocity and Trajectory Analysis

Cell Ranger (10X genomics) was used to process the FASTQ sequencing files, and its output was used to calculate RNA velocity using the package veloCyto ^63^. Seurat ^64^ was subsequently used to integrate the various RNA velocity output files and processed expression matrices. Scanpy ^65^ was used to process expression matrices for downstream analyses in python. Further RNA velocity was performed using scVelo ^66^, and dynamo ^67^ was used to visualise velocity vector fields. Initial trajectory analysis was performed using the CellRank package ^68^ using the RNA velocity kernel calculated using scVelo and the optimum transport kernel calculated using Moscot ^69^. The cell most highly associated with the inferred initial state of the velocity kernel was used as the initial cell for pseudotime analysis using Palantir after imputation ^70^. Where the kernels were used for inference of macrostates, the time-predominant kernel was calculated by linear combination of the optimum transport kernel, and connectivity kernel at ratio 0.8:0.2 to achieve optimum stability and emphasise time until sample collection as the major driving factor of cellular trajectory. The velocity-predominant kernel was calculated by linear combination of the velocity kernel and connectivity kernel at ratio 0.9:0.1.

### Perturbation analysis

Perturbation analysis was performed after filtering for the top 3000 highly variable genes based on the CINEMA-OT model ^29^ using shared monocytes vs haematoma-enriched monocytes as conditions.

### E-distance calculations

The scGPT model^30^ pre-trained on all human cell types was used to embed our data after filtering for the top 3000 highly variable genes using the standard tutorial. Then, the scPerturb python package was used to estimate the E-distance between the embeddings for the different cell types, days, and tissue^72^.

### Temporal trajectory analysis

Based on the scGPT embeddings of our data, we used scanpy’s diffusion tool to create a new pseudotime based on a random cell at the earliest timepoint. Generalized linear models were fitted in this trajectory to estimate the most important gene sets which are either up or downregulated. For each pathway database, genes whose coefficient had a p-value of less than 0.05 after Bonferroni correction were kept for the gene set enrichment analysis.

### Statistical Analysis

Statistical tests comparing hematoma and tissue were performed using the Wilcoxon test for paired samples with a p-value of 0.05 as a cutoff for significance.

## Data Availability

Raw sequencing files and metadata will be made available on GEO on publication. Codes used for analysis is available on GitHub (the link will be live on publication).

## Acknowledgements

National Institutes of Health grant R21NS108060 (LHS)

National Institutes of Health grant R01NS097728 (LHS)

American Heart Association 19EIA34770133 (LHS)

National Institutes of Health grant 1S10OD030363-01A1 (core facility)

## Author contributions

Conceptualization: YK, CJ, MA, LHS

Methodology: YK, LC, LHS

Investigation: YK, CJ, LC, SEV, JD, HB, CM, RH, BJC

Funding acquisition: LHS

Supervision: MA, LHS

Writing – original draft: YK, CJ, LC, JD

Writing – review & editing: YK, CJ, LC, MT, MA, LHS

## List of Supplementary Materials

Supplementary materials

Fig S1 to S15

Table S1

References (62*-80*)

