## Supplementary Material for "Single-Cell Analysis of Microglia and Monocyte Dynamics Uncover Distinct TNF-α-driven Neuroimmune Signatures after Intracerebral Hemorrhage"

**Single-Cell Temporal Atlas of Myeloid Cells in the Live Haemorrhagic Brain**

Supplemental Methods

**Sample processing**

Evacuated haematoma samples were stored in collection cups. Time-matched peripheral blood samples were collected perioperatively and stored in acid citrate dextrose (ACD) tubes. Time of collection, age, sex, type of surgery, and other clinically relevant variables were recorded and are reported here (Table S1). Samples were stored overnight at 4ºC prior to processing, which has previously been shown to induce minimal transcriptional changes in monocytes and other immune populations compared to fresh control ^1^.

Haematoma and peripheral blood (PB) samples were processed using a detailed protocol to minimize potential processing artifacts and batch effects in the resultant dataset^1^. All steps were performed in a sterile tissue culture hood. Haematoma fluid (HF) was strained through 40 uM filters to remove pieces of clot and brain. The haematoma clot itself was repeatedly washed with ice-cold HBSS and passed through a filter until liquid was light pink to clear. PB and HF samples were spun 500g x 10m and resuspended at 50% hematocrit in HBSS. Exosome supernatant was spun 2000g x 10m, aliquoted, and frozen at -80ºC. Granulocyte cells were then depleted from samples using a Rosette-Sep kit (Stem Cell #15664); protocol performed on ice and according to manufacturer’s instructions. Granulocytes comprise the dominant cell population in similar samples, so depletion was performed to increase the relative proportion of the immune cells of interest. Red blood cell (RBC) lysis was then performed by adding 1:10 1X RBC lysis buffer (BD #555899) and incubating at room temperature for 10m, repeated 1-2 times until sample was clear. Supernatant was then aspirated, and samples were resuspended in 5 mL of a 23% Percoll solution in PBS, spun 500g x 15m, aspirated and washed once with 5 mL complete RPMI. Cells were then counted using a hemocytometer and resuspended in PBS + 0.04% BSA at 1x10^6^ cells/mL.

**Sequencing and alignment**

A target of 10,000 cells from each blood and haematoma sample were processed using 10X Genomics 3’ Single Cell v3.1 kits. Cell suspensions were loaded onto Chromium Next GEN Chip G for GEM generation and barcoding. Other reagents were prepared and loaded according to the manufacturer's protocol. Library generation and amplification were performed by the Yale Center for Genome Analysis (YCGA) according to the manufacturer’s protocol. Libraries were then sequenced by the YCGA using the Illumina NovaSeq 6000 machine. Target sequencing depth was set at 200 million reads per sample for all blood samples and most haematoma samples. A reduced depth of 100 million reads per sample was set for haematoma samples with lower cDNA concentrations, since these tended to have substantially fewer cells. All samples were sequenced to a target sequencing saturation of 70 percent.

After sequencing, raw BCL files were converted to demultiplexed FASTQs using CellRanger software (3.1.0 for patients 1-8, 6.0.1 for patients 9-10). Samples were then aligned and gene expression matrices were generated using the CellRanger ‘count’ function, with default parameters except for an ‘expect-cells’ of 8,000. The default GRCh38 transcriptome (3.0.0) was used for alignment.

**Data preprocessing and integration**

All preprocessing was performed on the Yale Center for Research Computing’s high performance computing clusters. First, gene expression matrices were corrected for background expression using the SoupX R package (1.5.2) ^2^. For each sample, the background RNA profile (rho) was calculated using the AutoEstCont function and a default prior rho estimate of 0.05. Default CellRanger-generated clusters were used for the independent generation of rho estimates. The AdjustCounts function then adjusted the matrices to account for background contamination. Results were compared using known cell markers in brain and/or blood (*CD3A, S100A9, SPP1, MBP*). Lists of the genes most-often zeroed were checked to ensure legitimate cell types weren’t being removed. Adjusted matrices were exported to a Seurat object.

Subsequent data preprocessing was performed using R (4.0.5 and 4.3.3) and primarily the Seurat R package (4.0.0) ^3^. Cells with fewer than 200 features and features expressed in fewer than three cells were removed as a part of Seurat object generation. Percentage of mitochondrial genes was calculated using the percent.mito function. Quality control cutoffs were determined using covariate analysis of feature count (nFeature_RNA), total RNA count (nCount_RNA), and mitochondrial gene percentage for each cell. These cutoffs were refined based on downstream results of any likely doublet or dead cell clusters. Cutoffs were kept uniform within tissue type. For haematoma samples, cells with less than 20% mitochondrial RNA with between 600 and 6,500 features, with less than 35,000 RNA counts were selected. For blood samples, cells with less than 20% mitochondrial RNA with between 800 and 4,800 features, with less than 20,000 RNA counts were selected. Data was normalized using SCTransform, which uses a generalized linear model to regress out the technical confounder of sequencing depth within each cell. SCTransform has been shown to outperform other normalization methods when decoupling cell UMI count from expression for both highly expressed and lowly expressed genes ^4^.

Integration was performed using Seurat 3 ^5^ after a comparison between itself, Harmony, and LIGER. All three methods performed comparably on independent benchmarking analysis^6^. After comparing downstream clustering for all three, Seurat 3 was chosen for our analysis based on demonstrated removal of batch effects, minimizing clusters that split by patient rather than biological type. Specifically, reciprocal PCA (rPCA) integration was performed following normalization. The rPCA method is more conservative than other integration methods. It is recommended when samples have significant proportions of cell types that do not overlap (i.e. between blood and haematoma) and when there are many samples to integrate. Samples were integrated using 50 PCA dimensions and a k.anchor parameter of 20. We excluded lymphocytes, oligodendrocytes and astrocytes from our analysis to focus on myeloid cells for this study.

**Clustering**

Principal component cutoffs of the integrated assay were determined using ElbowPlot() and JackStraw() variance plots. Clustering was then performed with FindClusters() using the Louvain algorithm. For this and all downstream subclustering, the ‘resolution’ parameter of FindClusters() was iterated until differential expression analysis yielded the clearest split between defined cell types. For broad clustering, a resolution of 1.0 was ultimately used, identifying 33 clusters. Cluster annotation was manually performed through analysis of two sets of data. The first was output of differentially expressed genes generated using the limma implementation of the Wilcoxon Rank Sum test in the FindMarkers() function in Seurat, sorted by log-fold change. The second was comparison of relative expression levels of previously described lineage markers (*PTPRC, NKG7, CD3E, CSF1R, MS4A1, MBP*). Clusters were split into broad cell types (Myeloid, T/NK, CD45-/CNS, B Cells, or split populations).

Myeloid cells were then subclustered using FindClusters() at a resolution of 0.7. As with broad clustering, subcluster annotation was performed manually, by expression of canonical lineage markers and differentially expressed genes identified by FindMarkers(). Annotations were confirmed using SingleR v2.4.1 to map annotations from Hao et al. 2021 (for all populations) ^3^ and Mulder et al. 2021 (for myeloid populations) ^7^ onto this dataset. Doublets, dead cells, and non-myeloid cells were excluded from downstream analysis.

**Bulk RNA sequencing deconvolution**

Bulk RNA sequencing data was deconvoluted and aligned to our dataset using the protocol published by Marquez-Galera et al.^8^

**Cell type classification**

scTab^9^ was used to estimate the probabilities that a given cell in our data was classified as one of the hundreds of cell types available in the CellXGene data using the standard tutorial.

**Module scoring**

Gene sets for module scoring were created using lineage marker genes curated from previously published datasets ^10-12^. Module scores were calculated using the UCell package ^13^.

**Differential expression and gene set enrichment analysis**

Initial processing of single cell RNA sequencing data was performed on summed counts for each sample normalised using edgeR. Differential expression analysis was performed on these pseudobulk counts using limma ^60^ and compared using a generalised linear model for differences in gene expression by tissue, ICH volume, and GCS. Genes were ranked based on their T-statistic. Enrichment for good outcome was performed by comparing the difference in enrichment of genes between haematoma and peripheral blood samples separately for samples from patients with good outcome (mRS $\leq$ 3) and bad outcome (mRS > 3) after 3 months and comparing differentially regulated genes. These analyses were repeated with covariates incorporated into the limma formula as appropriate. Gene set enrichment analysis was performed using the MSigDB gene database for hallmark gene expression using the fgsea package ^61^ using a p-value of 0.05. Bulk RNA sequencing data were analysed in the same fashion but without the initial summation step. For enrichment analysis of gene lists without associated rank values, enrichR ^62^ was used with the MSigDB hallmark gene expression gene sets. The *EnhancedVolcano* and *ggplot2* packages were used for visualisation.

Supplementary Table 1. Patient Characteristics.

| **Age (years)** | **Sex** | **ICH location** | **ICH volume (mL)** | **IVH** | **GCS score** | **ICH score** | **Time from onset to surgery (hours)** | **Surgery type** |
| --- | --- | --- | --- | --- | --- | --- | --- | --- |
| 58 | Male | left frontotemporal | 84.8 | Yes | 15 | 2 | 62.0 | MIS |
| 58 | Male | left basal ganglia | 71.8 | Yes | 7 | 3 | 46.0 | MIS |
| 66 | Female | left parietal | 34.7 | Yes | 14 | 2 | 10.0 | MIS |
| 59 | Male | left basal ganglia | 35.8 | Yes | 13 | 1 | 91.0 | MIS |
| 71 | Male | right frontal | 60.7 | Yes | 8 | 3 | 11.3 | Craniotomy |
| 60 | Male | left cerebellar | 16.3 | No | 15 | 1 | 31.0 | Craniotomy |
| 70 | Female | right frontal | 57.0 | Yes | 13 | 2 | 48.3 | MIS |
| 55 | Female | right basal ganglia | 60.0 | Yes | 7 | 3 | 5.7 | Craniotomy |
| 68 | Male | left frontal | 42.6 | No | 15 | 1 | 99.5 | Craniotomy |
| 51 | Female | left basal ganglia | 81.9 | Yes | 10 | 3 | 290.5 | MIS |

ICH: intracerebral hemorrhage. IVH: intraventricular hemorrhage. GCS: Glasgow coma scale. MIS: minimally invasive surgery.

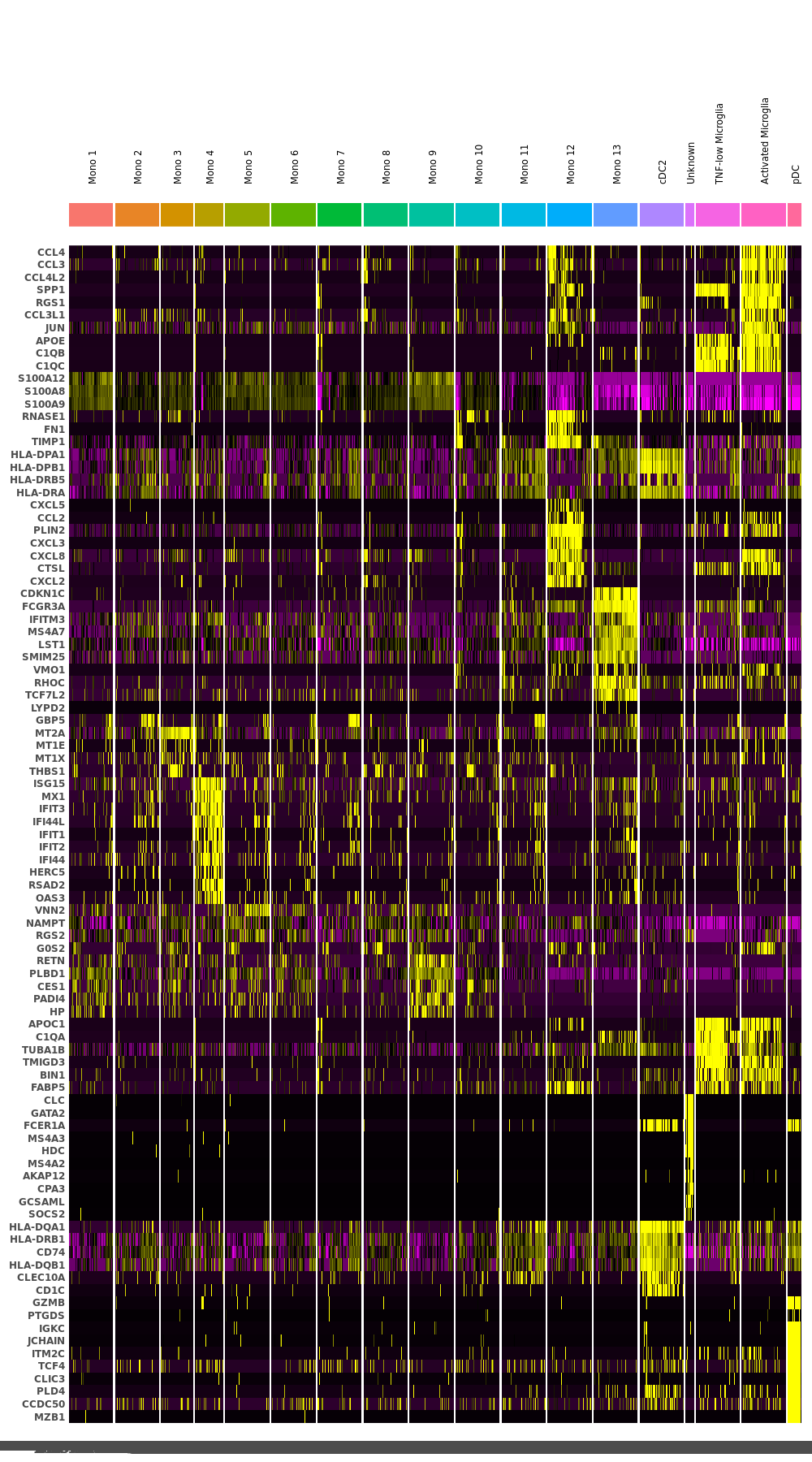

Supplementary Fig. 1. Heatmap of top 10 marker genes for each myeloid subcluster.

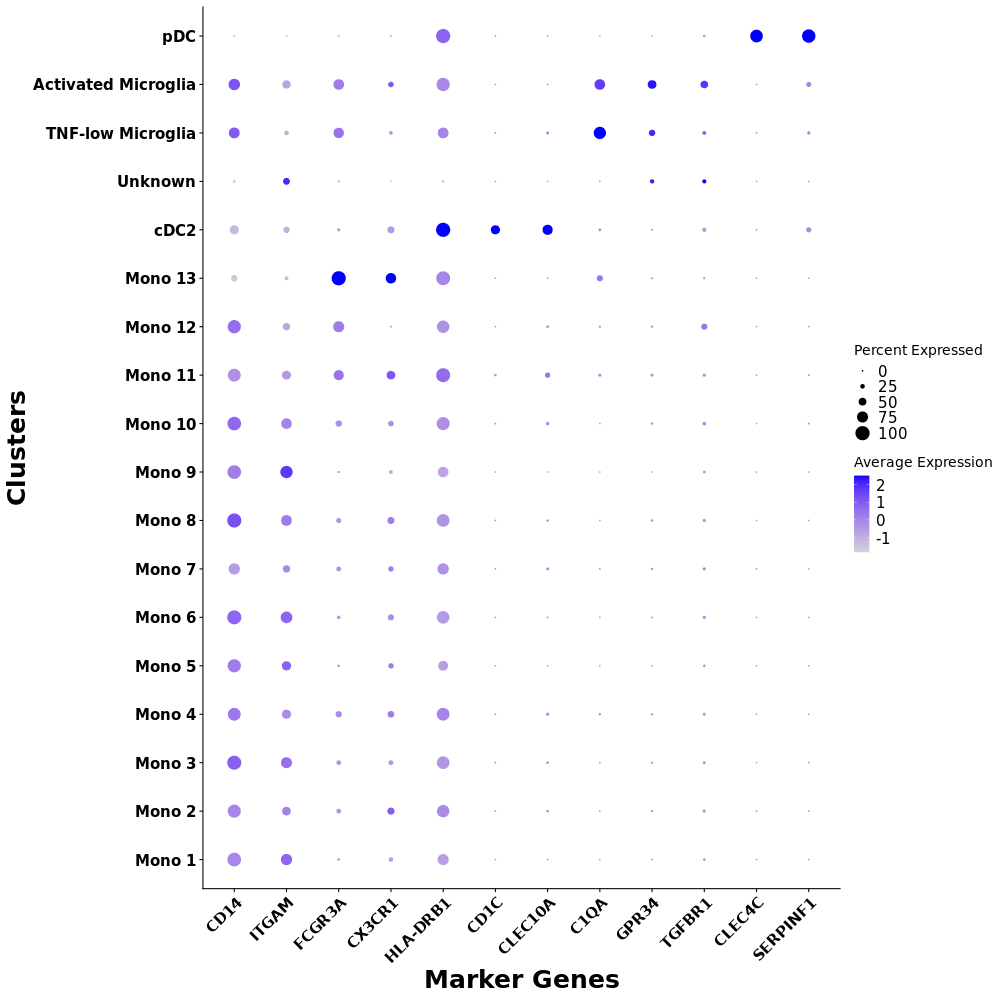

Supplementary Fig. 2. Dot plots demonstrating selected lineage marker expression in each cluster.

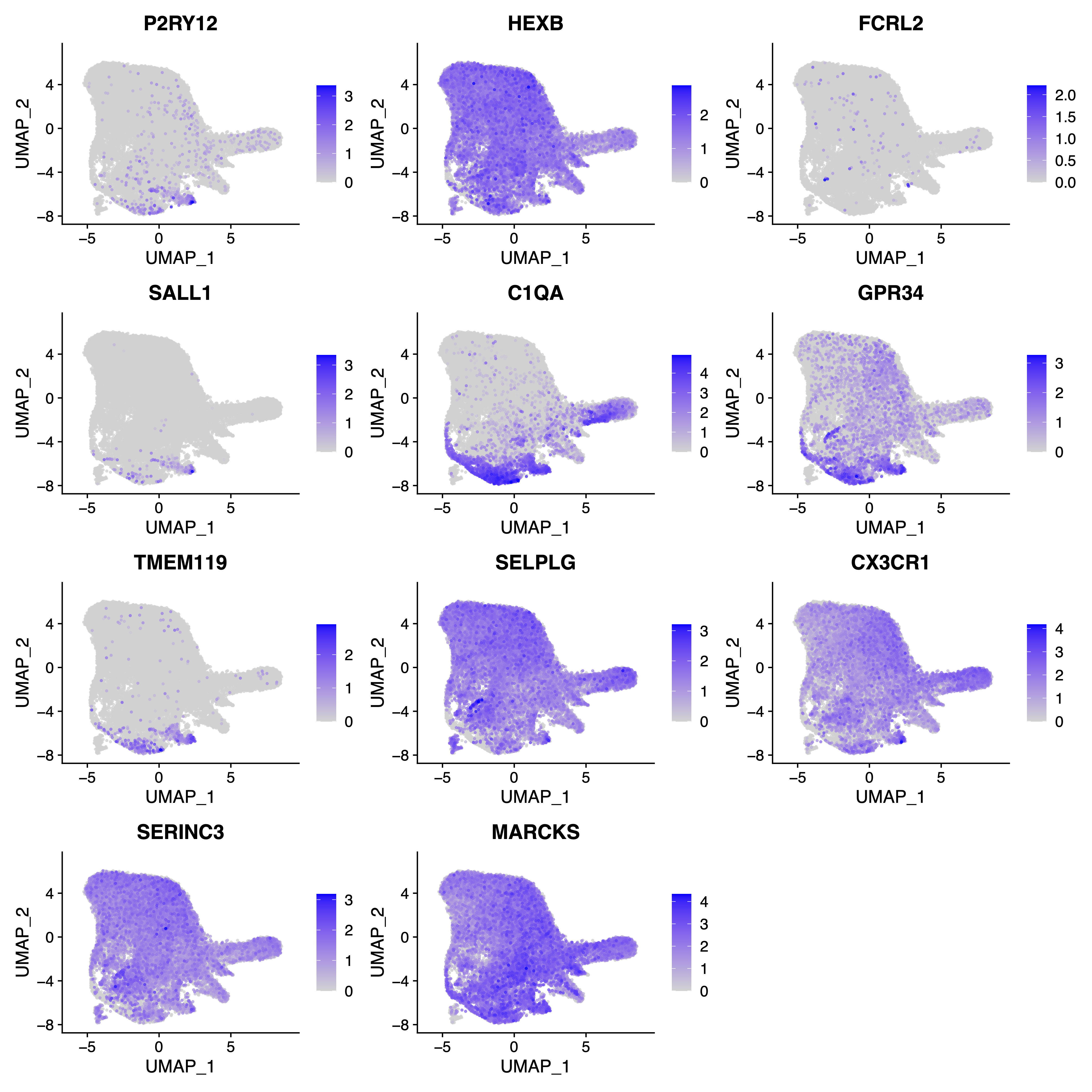

Supplementary Fig. 3. UMAP plots demonstrating expression of an extended array of marker genes for homeostatic microglia.

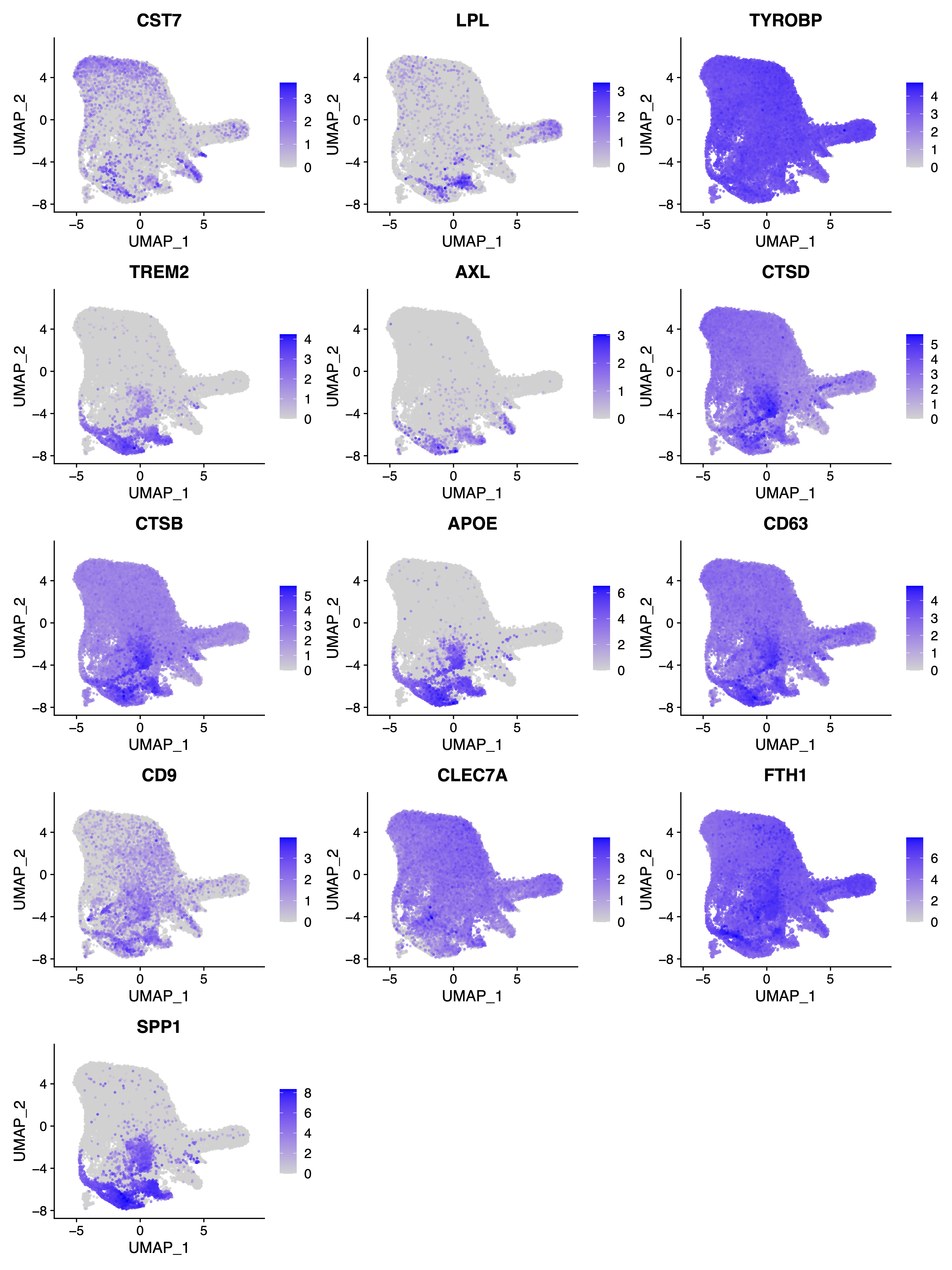

Supplementary Fig. 4. UMAP plots demonstrating expression of marker genes for disease-associated microglia.

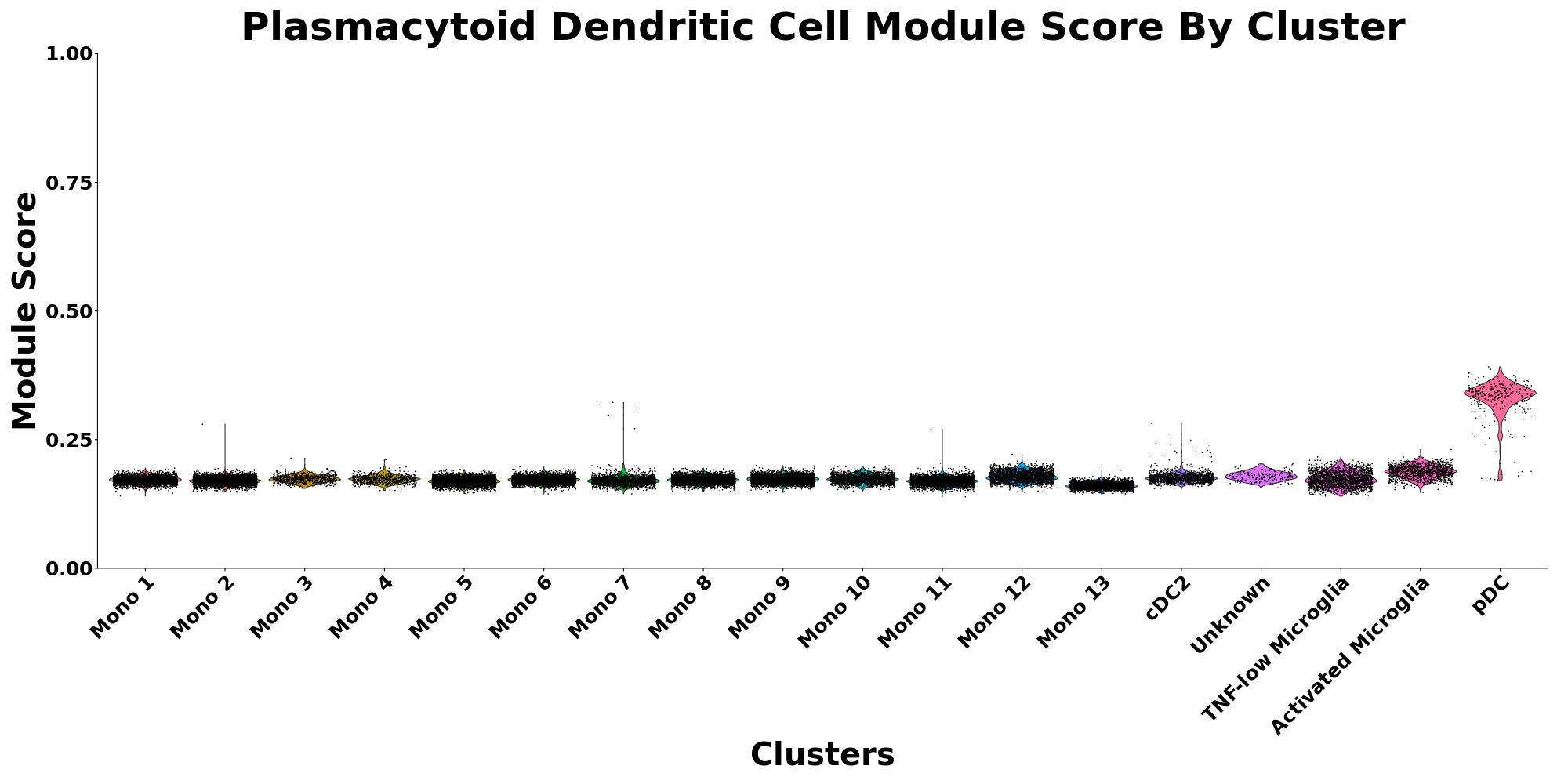

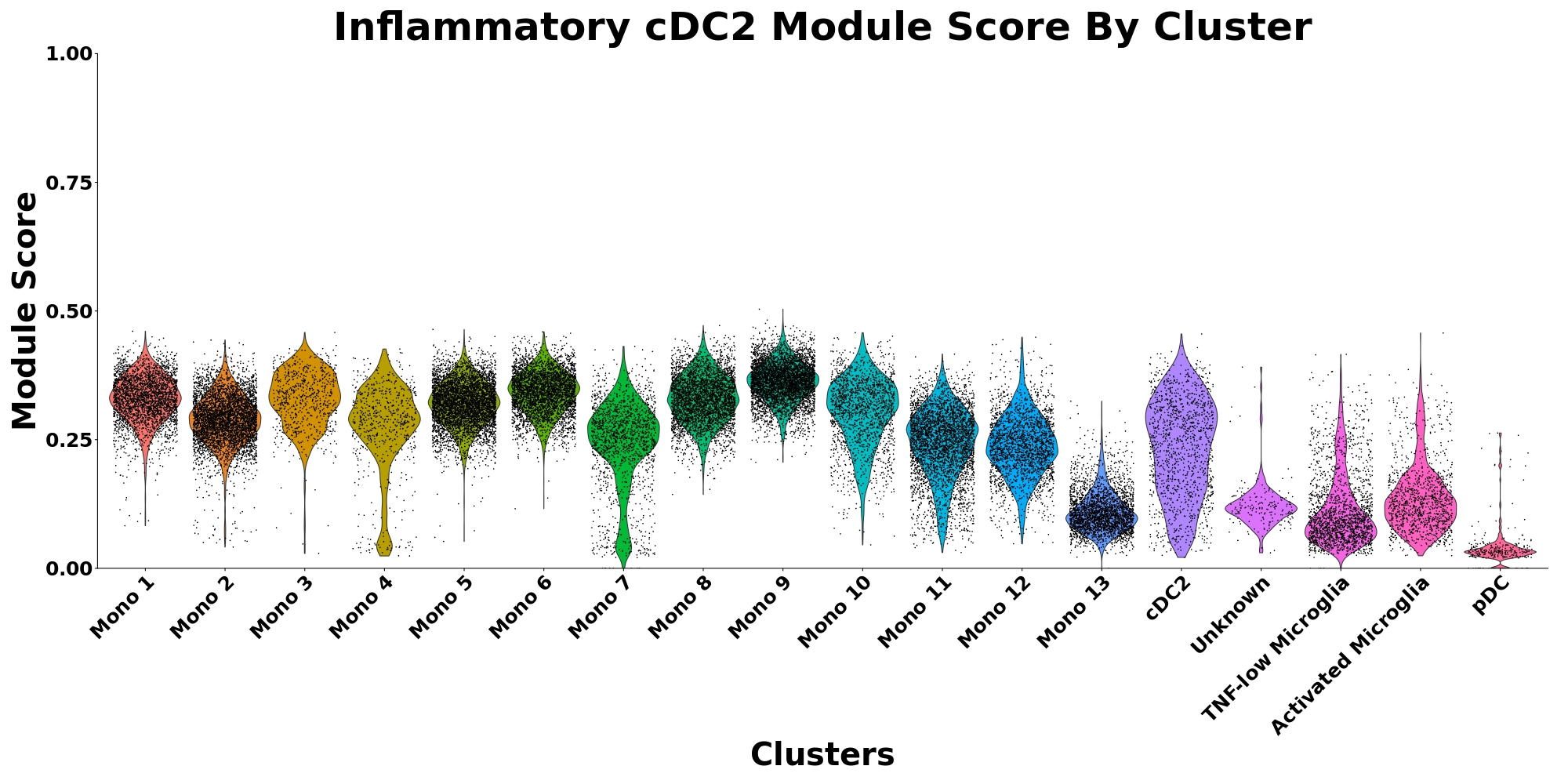

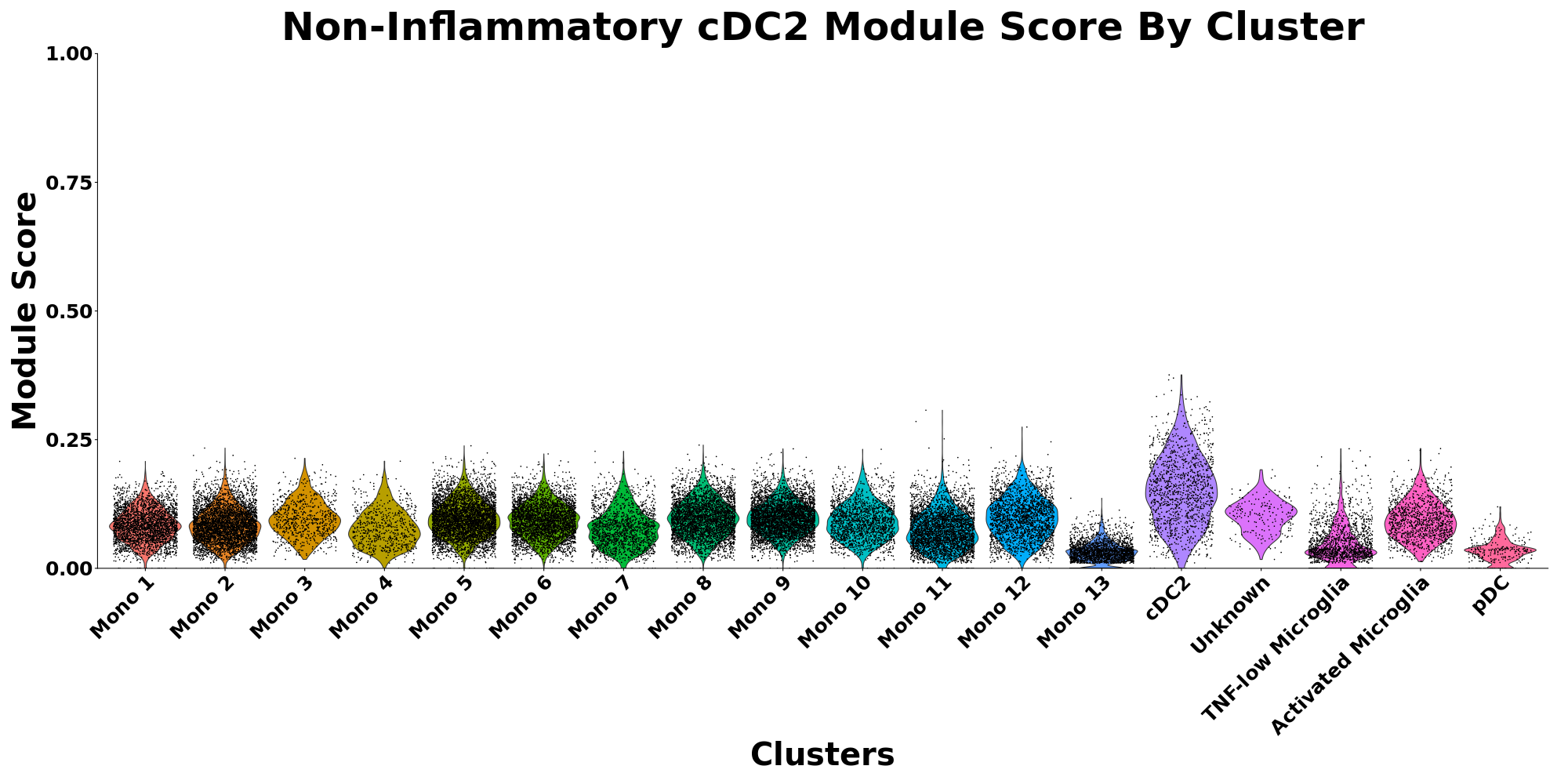

Supplementary Fig. 5. Module score plots plasmacytoid dendritic cells, inflammatory, and non-inflammatory cDC2 marker genes.

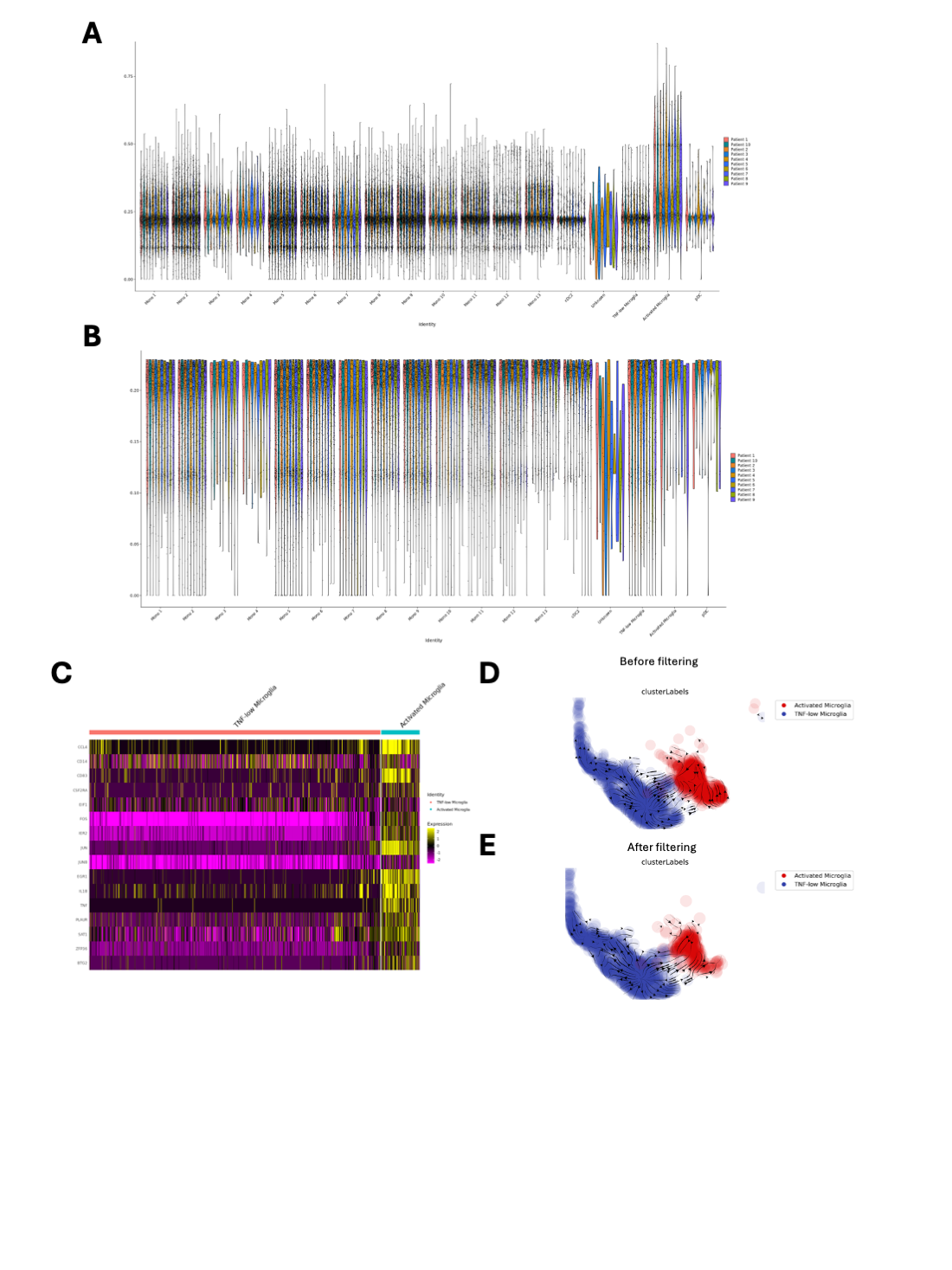

Supplementary Fig. 6. A. Module score plots for genes associated with damaged microglia excluding TNF pathway genes before filtering. B. Module score plots for genes associated with damaged microglia excluding TNF pathway genes after filtering for cells with low scores. C. Heatmap of DIM markers after filtering for low damage scores. D. RNA streamline plot for microglia before and after filtering for low damage scores.

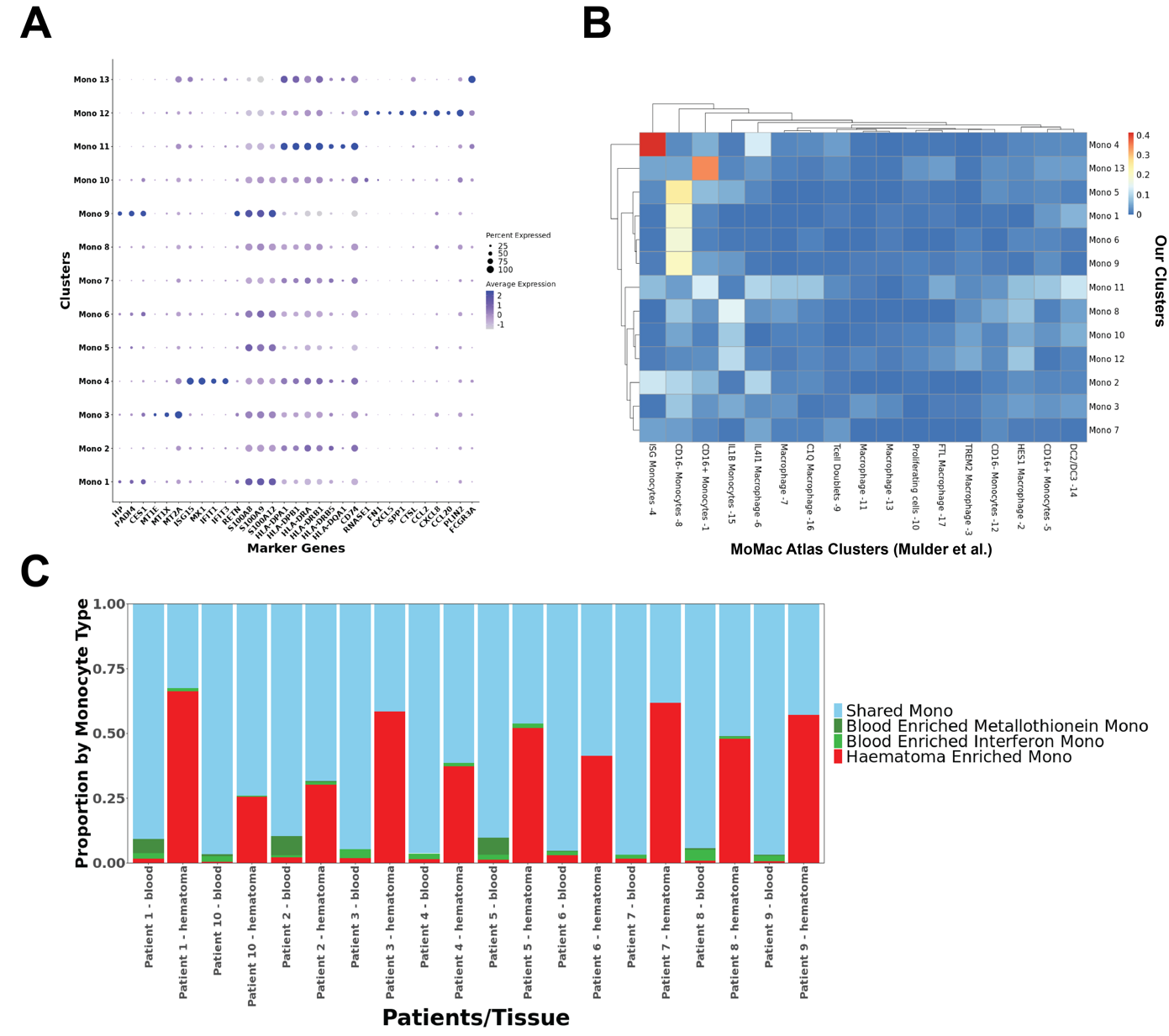

Supplementary Fig. 7. A. Dot plot showing top marker genes for CD14^+^ monocytes. B. Comparison of marker genes of monocytes in our dataset and a cross-tissue monocyte/microglia atlas^7^. C. Proportion plot by patient, by tissue.

Note, clusters 1-2 and 5-11 demonstrated relatively high levels of classical CD14^+^ monocyte markers and expressed high levels of genes such as *RETN*, *S100A8*, *S100A9*, *S100A12*, *PLBD,* and *NAMPT* (Fig. S1), which are found in peripheral blood after acute myocardial infarction and associated with worse left ventricular function due to increased infiltration^14^. These clusters were associated with haem metabolism and reactive oxygen species (Fig. 3A), further suggesting that these subsets could be activated monocytes which form part of the systemic response to ischaemia.

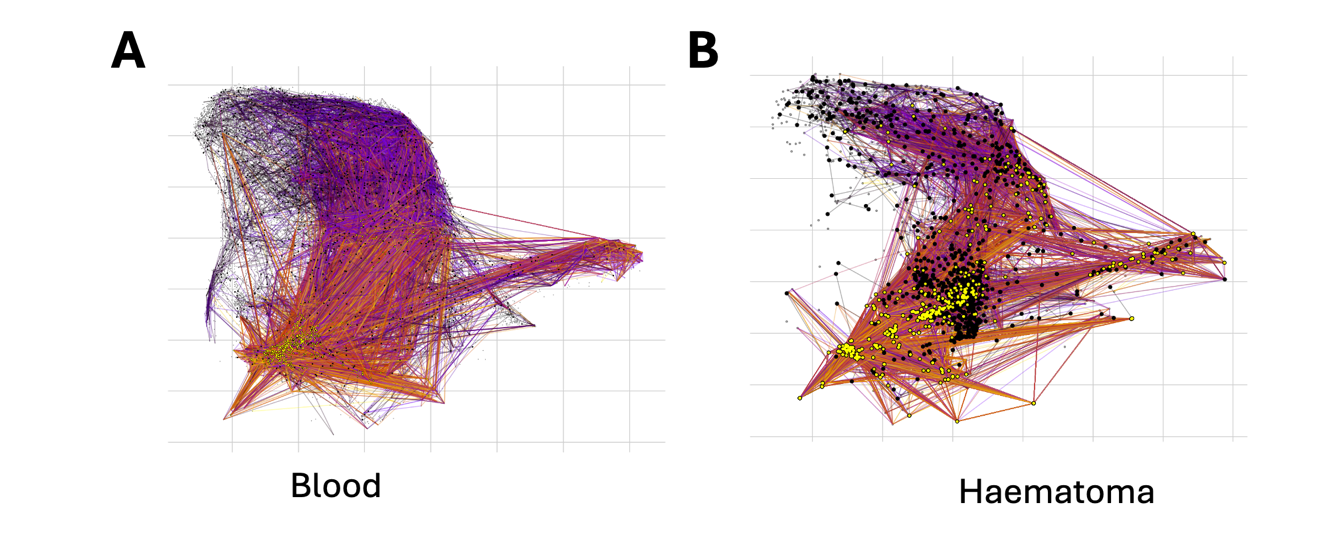

Supplementary Fig. 8. Random walk plots for peripheral blood and haematoma samples. **A:** Random walk plots of peripheral blood samples based on velocity (Black dots represent starting points, and red dot end points of random walks.) **B:** Random walk plots of haematoma samples based on velocity kernel

Supplementary Fig. 9. scGPT temporal embeddings. A. UMAP of model embedding coloured by days post injury. B. Energy distance plot from samples at earliest time point for Mono 12 cells in blood.

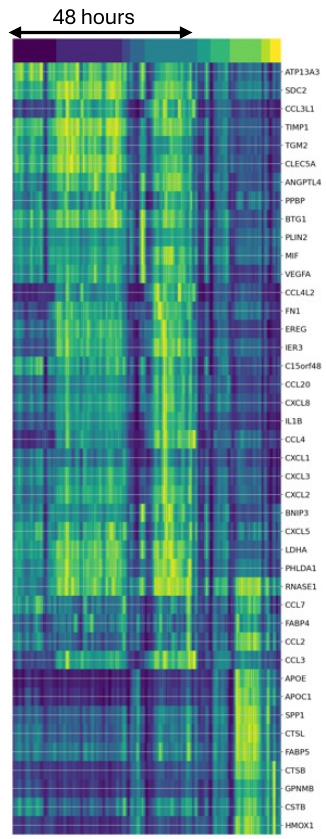

Supplementary Fig. 10. Inferred transcription rate of genes associated with the TNF-a/NFkB pathway. Individual cells are ordered by day after sample collection (increasing towards the right), and coloured by transcription rate (yellow: high, blue: low)

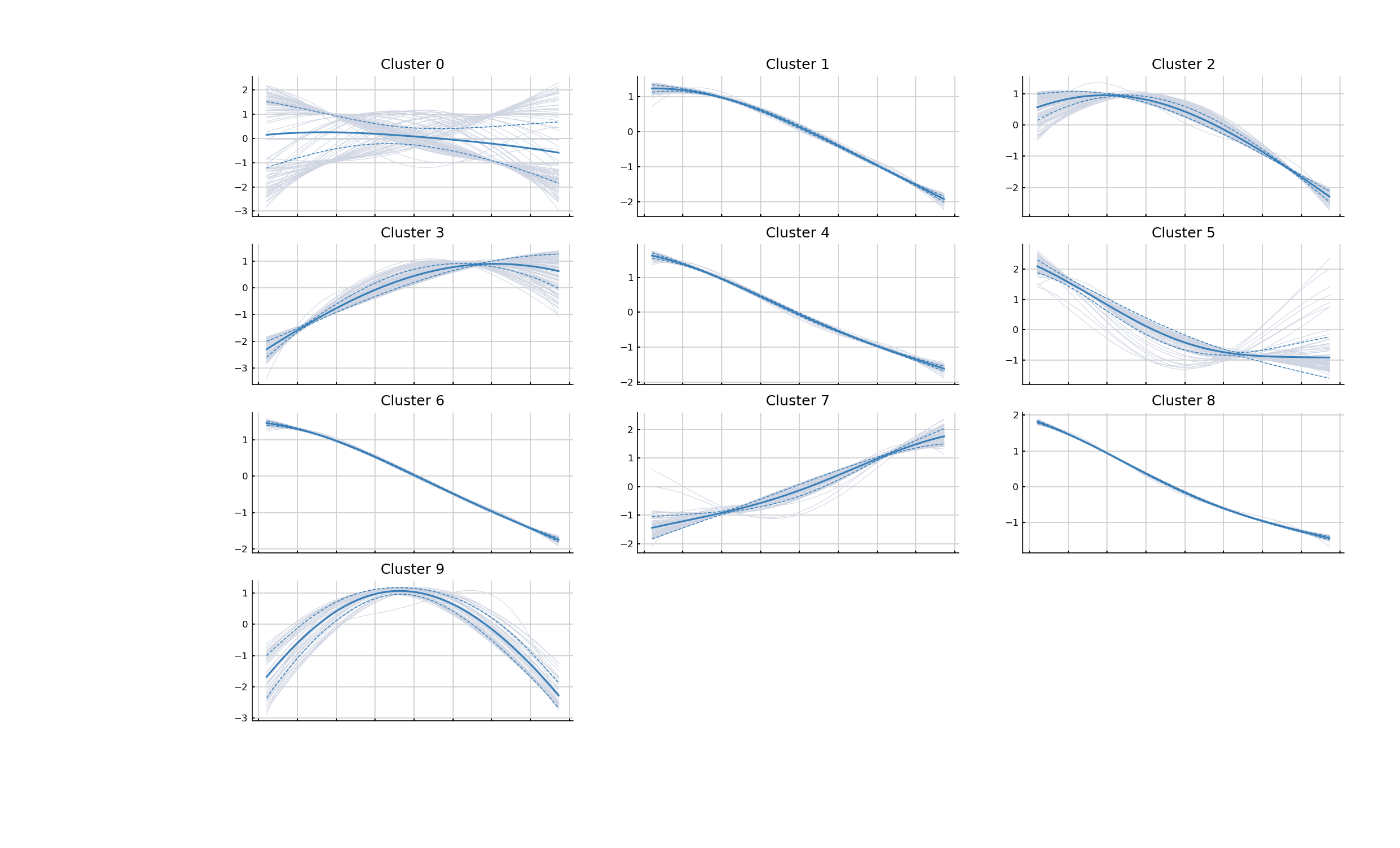

**A**

**B**

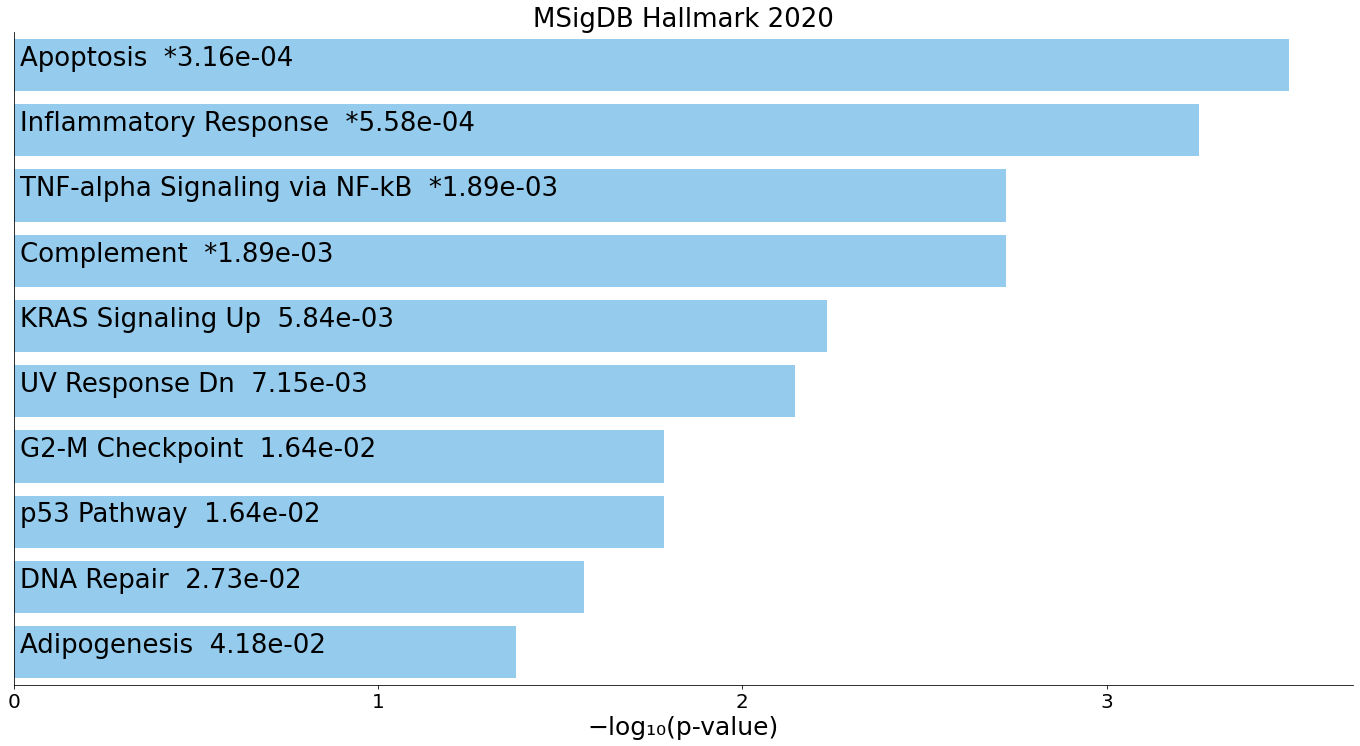

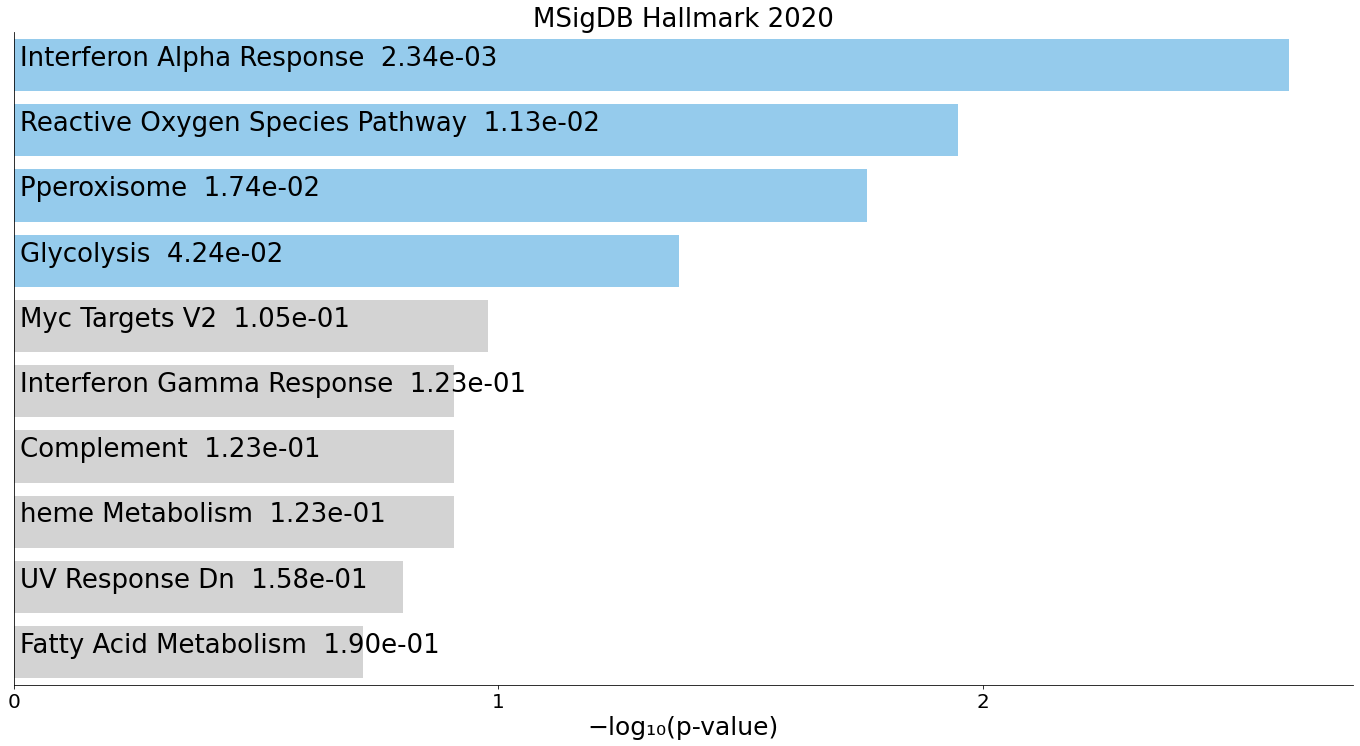

**C**

Supplementary Fig. 11. Gene expression patterns in Mono 12 cluster. **A**. Plots showing trends of various clusters of genes identified by similar behaviour over time. **B.** Gene sets of genes which exhibited monotone increase with optimum transport trajectory analysis (Cluster 3, 7) **C.** Gene sets of genes which exhibited monotone increase with optimum transport trajectory analysis (Clusters 1, 4, 6, 8)

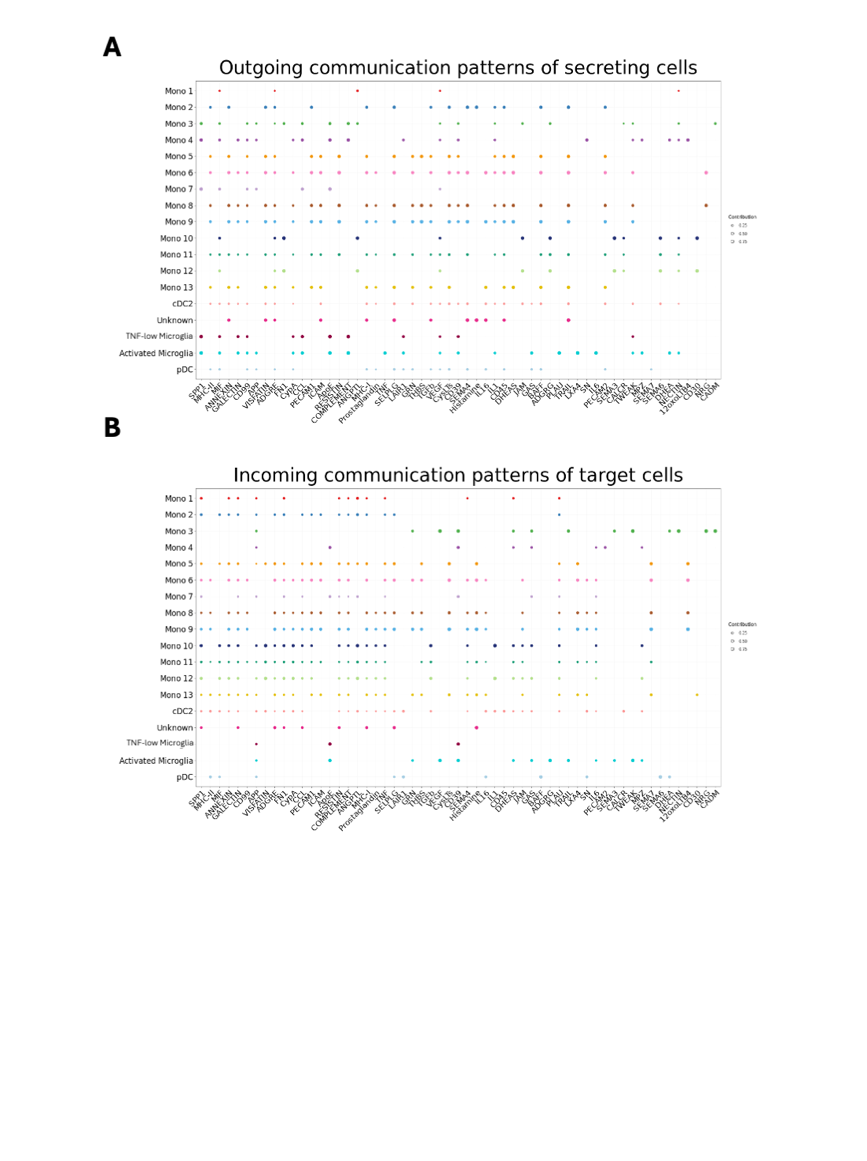

Supplementary Fig. 12. Patterns of ligand-receptor signalling in myeloid cells. A. Outgoing patterns and B. incoming patterns of ligand-receptor signalling.

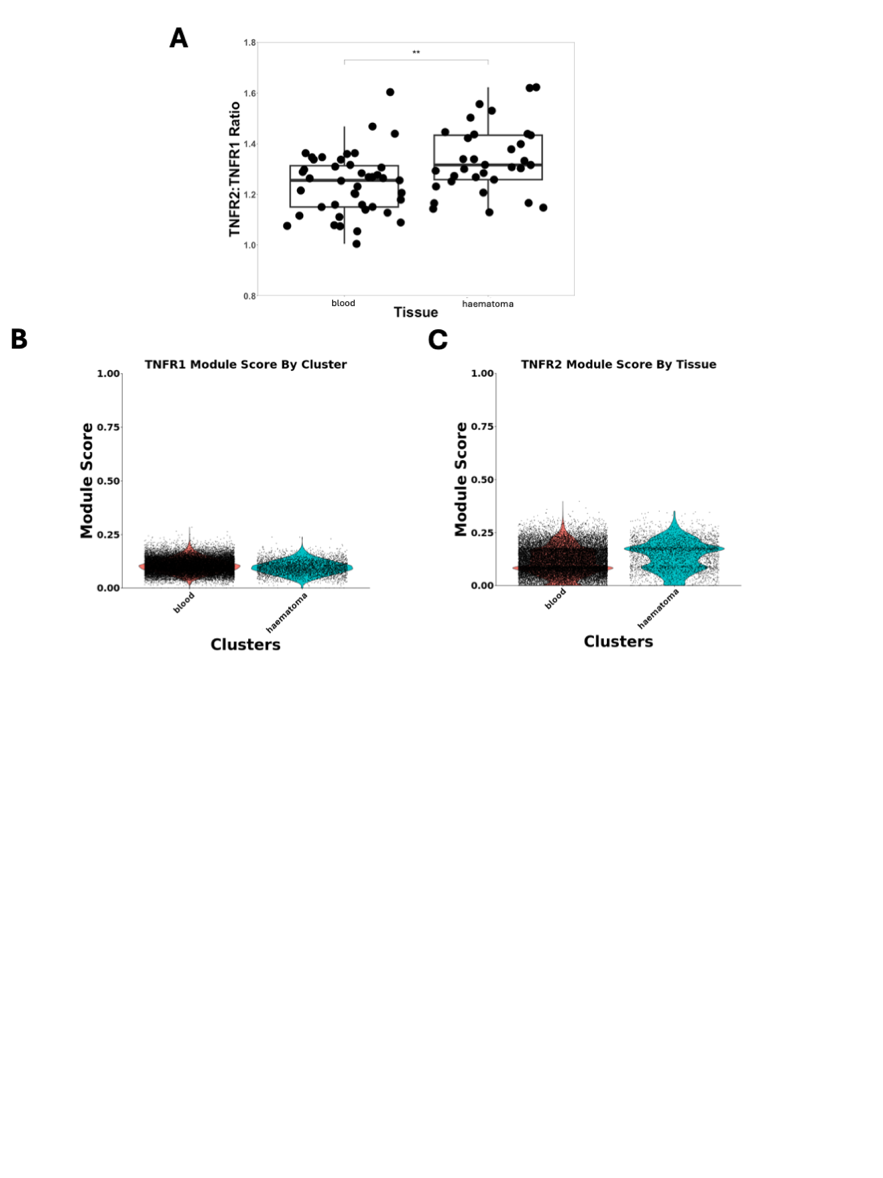

Supplementary Fig. 13. *TNFR1* and *TNFR2* expression in CD14+ monocytes. **A**. Boxplot of *TNFR2*:*TNFR1* transcription in bulk sequenced CD14+ monocyte samples from the MISTIE III trial. **B.** Module score for *TNFR1* signalling by tissue in pseudobulked CD14 monocytes (our cohort). **C.** Module score for *TNFR2* signalling by tissue in pseudobulked CD14 monocytes (our cohort). **: p<0.01, Wilcoxon test

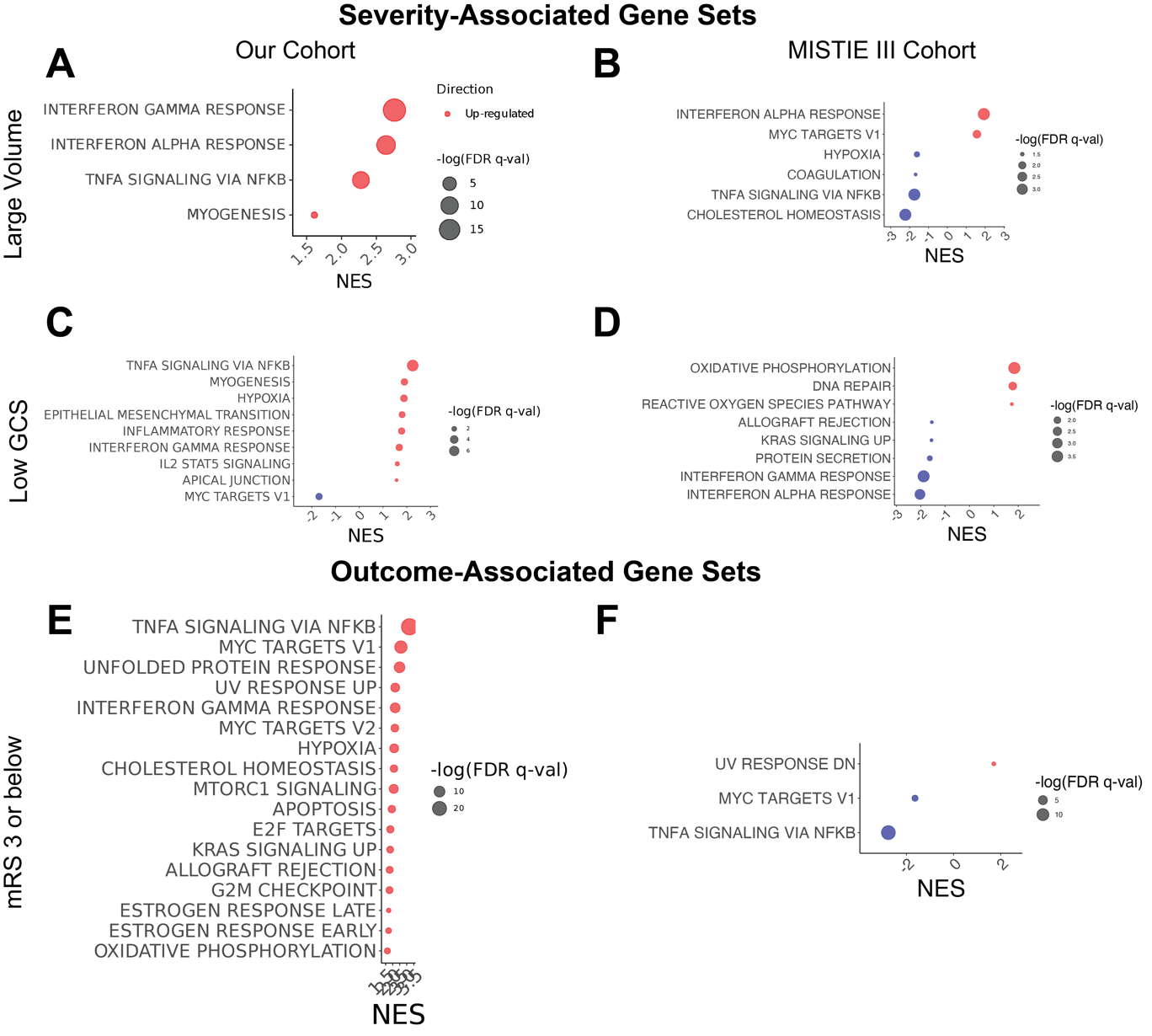

Supplementary Fig. 14. Gene signatures associated with severity and outcome in peripheral blood.

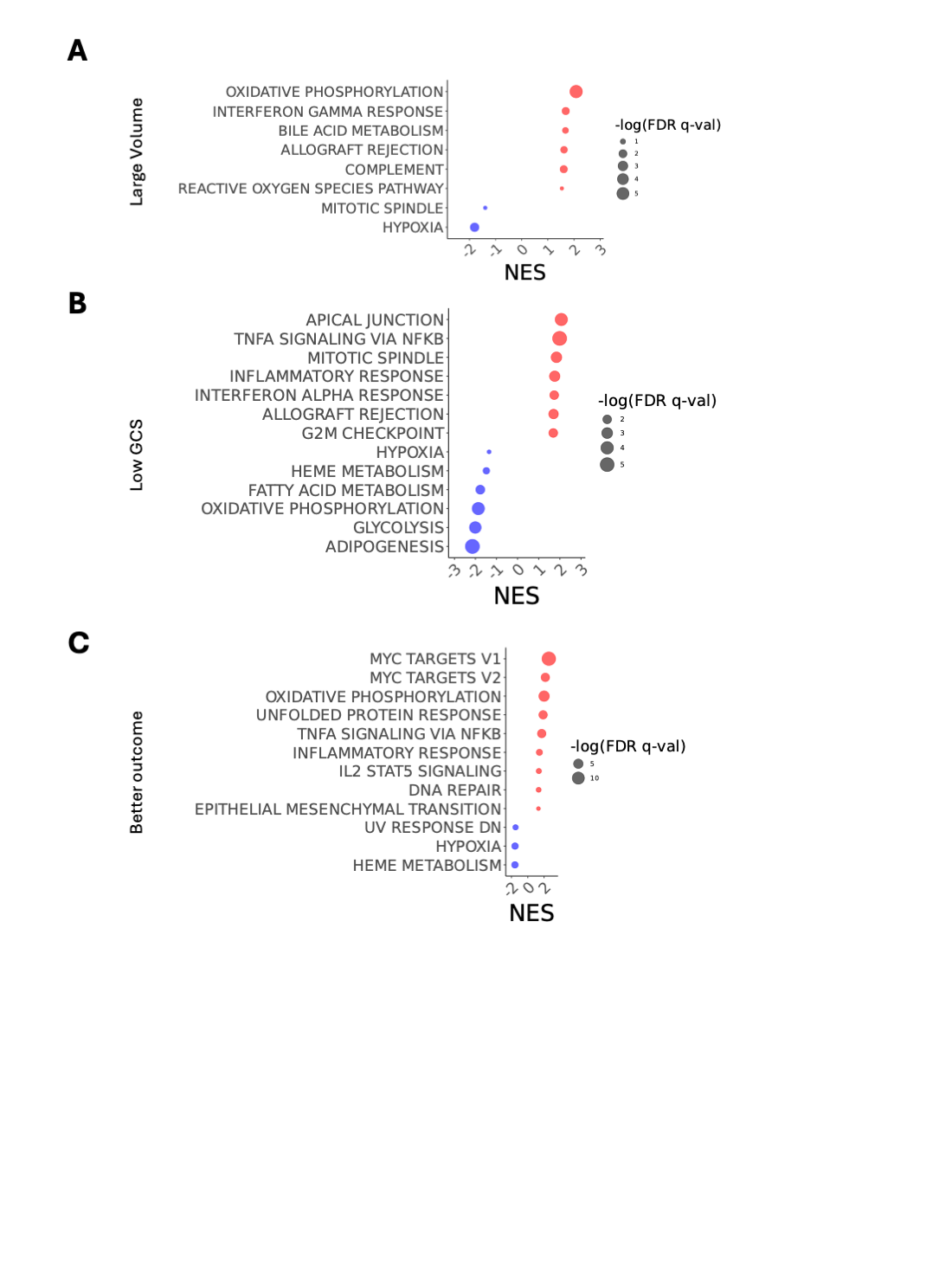

Supplementary Fig. 15. Gene signatures associated with severity and outcome in Mono 12.
